# A Modified TurboID Approach Identifies Tissue-Specific Centriolar Components In *C. elegans*

**DOI:** 10.1101/2021.12.20.473533

**Authors:** Elisabeth Holzer, Cornelia Rumpf-Kienzl, Sebastian Falk, Alexander Dammermann

## Abstract

Proximity-dependent labeling approaches such as BioID have been a great boon to studies of protein-protein interactions in the context of cytoskeletal structures such as centrosomes which are poorly amenable to traditional biochemical approaches like immunoprecipitation and tandem affinity purification. Yet, these methods have so far not been applied extensively to invertebrate experimental models such as *C. elegans* given the long labeling times required for the original promiscuous biotin ligase variant BirA*. Here, we show that the recently developed variant TurboID successfully probes the interactomes of both stably associated (SPD-5) and dynamically localized (PLK-1) centrosomal components. We further develop an indirect proximity labeling method employing a GFP nanobody-TurboID fusion, which allows the identification of protein interactors in a tissue-specific manner in the context of the whole animal. Critically, this approach utilizes available endogenous GFP fusions, avoiding the need to generate multiple additional strains for each target protein and the potential complications associated with overexpressing the protein from transgenes. Using this method, we identify homologs of two highly conserved centriolar components, Cep97 and Bld10/Cep135, which are present in various somatic tissues of the worm. Surprisingly, neither protein is expressed in early embryos, likely explaining why these proteins have escaped attention until now. Our work expands the experimental repertoire for *C. elegans* and opens the door for further studies of tissue-specific variation in centrosome architecture.

## INTRODUCTION

Studies in the nematode *C. elegans* have contributed greatly to our understanding of centrosome biology, with comprehensive genome-wide screens leading to the identification of the conserved molecular machinery underlying centriole assembly as well as key players in mitotic pericentriolar material (PCM) recruitment, aided by the strong and reproducible phenotypes observed following RNAi-mediated depletion in the early embryo (Oegema & Hyman, 2006; Pintard & Bowerman, 2019). The striking success of biochemical approaches reconstituting elements of centriole and PCM assembly *in vitro* underline the power of this experimental model (Kitagawa *et al*, 2011; van Breugel *et al*, 2011; Woodruff *et al*, 2017; Woodruff *et al*, 2015). The assumption here is that we have a complete parts list of all essential components of centrosomal structures. Recent work identifying further centriolar and pericentriolar material components whose depletion phenotypes are somewhat more subtle (Erpf *et al*, 2019; Sugioka *et al*, 2017; von Tobel *et al*, 2014) suggests this is not necessarily the case. Furthermore, while most of the work in *C. elegans* has focused on the early embryo, the available data for the acentriolar centrosome at the ciliary base of sensory neurons (Garbrecht *et al*, 2021; Magescas *et al*, 2021) and the non-centrosomal microtubule organizing center in the intestine (Feldman & Priess, 2012; Sanchez *et al*, 2021) (Fig. 1A) hints at tissue-dependent differences in centrosome composition and assembly mechanisms in somatic tissues of the worm that as yet remain little explored.

**Figure 1:**
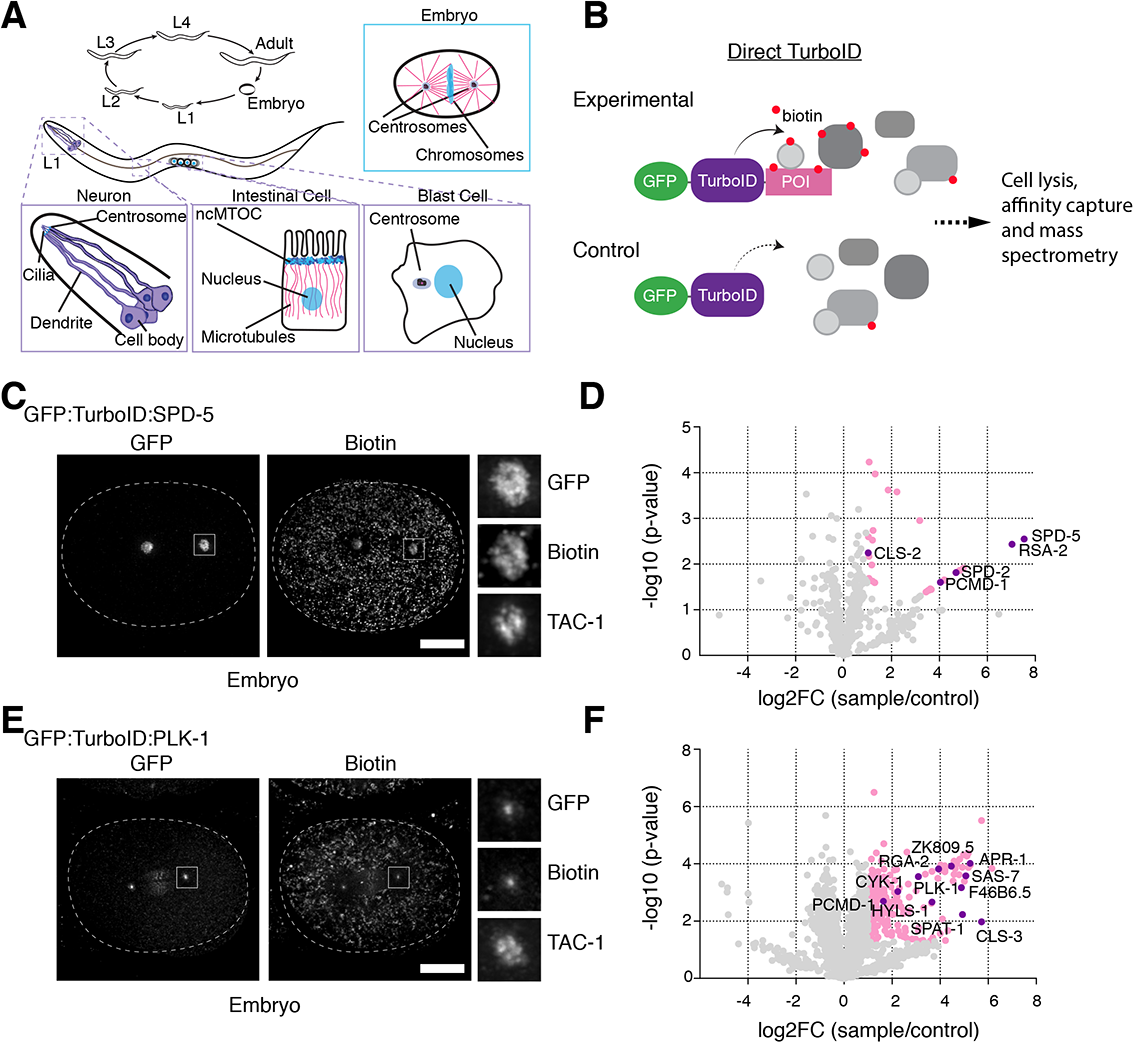
Probing the proximity interactome of centrosomal proteins by TurboID. (**A**) Centrosomes and centrosome-related structures in *C. elegans*. Centrosome assembly and function has been primarily examined in the early embryo, although some limited work has also been performed on the acentriolar centrosome at the ciliary base of sensory neurons (Garbrecht *et al.*, 2021; Magescas *et al.*, 2021) and the non-centrosomal microtubule organizing center in the intestine (Feldman & Priess, 2012; Sanchez *et al.*, 2021). Blast cells are two pairs of cells in the L1 larva that will give rise to the somatic gonad and germline in the adult worm. At the L1 larval stage these cells are arrested in G1 and G2 phase of the cell cycle, respectively, with what appear to be canonical interphase centrosomes (Fukuyama *et al.*, 2006; Hong *et al.*, 1998). (**B**) Schematic of direct TurboID (Branon *et al.*, 2018) which in our implementation also includes a GFP tag to visualize the TurboID fusion. Note that experimental and control TurboID fusions (lacking the protein of interest, POI) are expressed as separate transgenes, though under the same promoter and 3’ regulatory sequences. (**C**) TurboID applied to the PCM scaffolding protein SPD-5. Immunofluorescence micrograph of early embryo expressing GFP:TurboID:SPD-5 stained for GFP, biotin (streptavidin) and TAC-1 as a PCM countermarker. Biotinylation signal is observed at centrosomes without supplemental biotin addition. (**D**) Result of LC-MS/MS analysis for direct TurboID on SPD-5 from mixed-stage embryos. Volcano plot of −log10 p-values against log2 fold change (sample/control). Significantly enriched proteins (Log2 enrichment >1, p-value <0.05) are indicated in pink, with selected proteins highlighted. See also Supplemental Table 1 and Fig. S2C. (**E**) TurboID applied to the dynamically localized PCM regulator PLK-1. Immunofluorescence micrograph of early embryo expressing GFP:TurboID:PLK1 stained for GFP, biotin (streptavidin) and TAC-1 as a PCM countermarker. Biotinylation signal is observed at centrioles coincident with GFP:PLK-1 signal. (**D**) Result of LC-MS/MS analysis for direct TurboID on PLK-1 from mixed-stage embryos. See also Supplemental Table 1 and Fig. S2D. Scale bars in C and E are 10µm.

Interaction proteomics potentially offers an alternative strategy to identifying novel components of centrioles and centrosomes in different tissues of the worm. However, traditional approaches to probing cytoskeletal protein-protein interactions have been hampered by the fact that these interactions mostly take place in the context of the fully assembled structure, with few soluble cytoplasmic pre-complexes available for isolation by immunoprecipitation or other biochemical methods (Wueseke *et al*, 2014). Proximity-dependent methods circumvent this problem by direct labeling of interacting/proximal proteins, eliminating the need to preserve protein-protein interactions during extract preparation (Roux *et al*, 2012). BioID has revolutionized vertebrate centrosome biology (Firat-Karalar *et al*, 2014; Gheiratmand *et al*, 2019; Gupta *et al*, 2015) but hitherto has not been applied to worms (or flies) due to the low activity of the original promiscuous biotin ligase BirA* requiring long labeling times (∼24 hours) incompatible with the rapid development of invertebrate model organisms. The variant TurboID reduces this time to as little as 10 min and indeed this tool was developed specifically for use in *C. elegans* and *Drosophila* (Branon *et al*, 2018). Here, we examine the potential for TurboID to probe the interactomes of both stably associated and dynamically localized centrosomal components in *C. elegans* embryos. We further develop an indirect proximity labeling method and use it to gain initial insights into the tissue-specific variation in centrosome composition in the worm, identifying several proteins expressed exclusively at later developmental stages, including potential homologs of two highly conserved centriolar components, Cep97 and Bld10/Cep135.

## RESULTS AND DISCUSSION

### Establishment of TurboID proximity-dependent labeling to probe centrosome architecture in *C. elegans*

When we set out on this project TurboID had not yet been successfully applied to identify the proximity interactors of specific proteins in *C. elegans* (two such studies were recently published by the Feldman and de Bono labs, (Artan *et al*, 2021; Sanchez *et al.*, 2021)). To explore the potential for TurboID to map centrosomal protein-protein interactions we chose two target proteins for our proof of principle experiments, the PCM scaffolding protein SPD-5 and the Polo-like kinase PLK-1. As the major structural component of centrosomes in *C. elegans* (Hamill *et al*, 2002; Woodruff *et al.*, 2015), SPD-5 displays no detectable exchange with the cytoplasmic pool once incorporated into the centrosome (Laos *et al*, 2015). Moreover, fluorescence correlation spectroscopy has shown that SPD-5 is largely monomeric in the cytoplasm, with only a small fraction associated with the PP2A regulatory proteins RSA-1 and RSA-2 (Wueseke *et al.*, 2014). Consistent with this, our efforts to identify SPD-5 interactors by tandem affinity purification using the localization and affinity purification (LAP) tag (Cheeseman & Desai, 2005) were unsuccessful (our unpublished observations). In contrast, Plk1 dynamically localizes to multiple mitotic structures, with almost complete exchange at centrosomes within seconds after photobleaching. A key regulator of mitotic events, its behavior is thought to be the result of transient interactions with its numerous substrates throughout the cell (Kishi *et al*, 2009; Mahen *et al*, 2011). SPD-5 and PLK-1 both therefore in different ways represent challenging subjects for interaction biochemistry. We began by generating transgenic strains for each protein expressing a GFP-TurboID fusion under endogenous regulatory sequences by Mos1 transposon mediated insertion at a defined chromosomal locus (Frokjaer-Jensen *et al*, 2008), along with corresponding control strains lacking the SPD-5/PLK-1 coding sequence (Fig. 1B). Both fusions localized similarly to the endogenous protein in early embryos, with GFP:TurboID:SPD-5 distributed throughout the PCM while GFP:TurboID:PLK-1 was heavily concentrated at centrioles as well as at other cellular locations, including kinetochores and the spindle midzone (Fig. 1C, E). Fluorescent streptavidin probes showed biotinylation signal coincident with GFP fluorescence in the same locations, indicating functionality of the biotin ligase. In contrast there was little biotinylation signal in control strains beyond weak mitochondrial staining also seen in N2 wild-type embryos (not shown). Interestingly, biotinylation staining, both specific and non-specific, was not noticeably increased upon addition of supplemental biotin, nor were there any deleterious consequences to growing TurboID strains on biotin-producing OP50 bacteria (Embryonic viability 99.4%, n=1890, GFP:TurboID:SPD-5; 99.9%, n=2622 GFP:TurboID:PLK-1). We therefore performed *C. elegans* liquid cultures and embryo isolations under standard conditions, with three independent replicates for both experimental (GFP:TurboID:SPD-5/PLK-1) and control (GFP:TurboID) conditions.

In order to enrich for centrosomal interactors, crude embryo extracts were separated into soluble (cytoplasmic) and insoluble (cytoskeletal) fractions, the latter containing the centrosomes as confirmed by characteristic foci of GFP signal when examined under the fluorescence microscope. Centrosomal pellets were then resolubilized under denaturing conditions and used for streptavidin pulldowns followed by LC-MS/MS analysis. As also noted by Artan et al.(Artan *et al.*, 2021) endogenously biotinylated carboxylases represented a significant proportion of the isolated peptides. However, they, along with other common contaminants including vitellogenins and mitochondrial proteins, are largely filtered out by considering only those proteins significantly enriched in experimental samples over controls (Log2 enrichment >1, p-value <0.05). What remains for SPD-5 is a remarkably short list, 30 proteins (Fig. 1D, Supplemental Table 1). By far the most enriched protein is SPD-5 itself, along with its interacting partner RSA-2, though somewhat surprisingly not the PP2A complex of RSA-1, PAA-1 and LET-92 with which SPD-5 has been reported to interact in the cytoplasm via RSA-2 (Schlaitz *et al*, 2007), none of which were detected in the prep. Also highly enriched are SPD-2, which contributes to SPD-5 self-assembly *in vitro* (Woodruff *et al.*, 2015) and centrosome maturation *in vivo* (Kemp *et al*, 2004; Pelletier *et al*, 2004), and the recently described PCM tethering factor PCMD-1 (Erpf *et al.*, 2019). The SPD-2 recruitment factor SAS-7 (Sugioka *et al.*, 2017), also highly enriched, just failed the significance threshold, while the remaining proteins are either uncharacterized or likely contaminants (Fig. 1D, Supplemental Table 1). The list of SPD-5 proximity interactors therefore consists primarily of proteins related to PCM scaffold assembly. Notably absent are other centriolar and PCM components, including highly abundant ones such as γ-tubulin (TBG-1 in *C. elegans*)(Bauer *et al*, 2016). TurboID, then, is highly specific with a labeling radius narrower than the 10-15 nm estimated for BirA*(Kim *et al*, 2014). The list for PLK-1 is considerably longer, 235 proteins, including numerous proteins involved in cytoskeletal organization and morphogenesis, such as CLS-3/CLASP(Espiritu *et al*, 2012), CYK-1(Swan *et al*, 1998), RGA-2(Diogon *et al*, 2007), APR-1/APC(Mizumoto & Sawa, 2007) and several uncharacterized proteins reported to interact with PLK-1 in high-throughput screens (F46B6.5, ZK809.5, (Boxem *et al*, 2008; Zhong & Sternberg, 2006)) (Fig. 1F, Supplemental Table 1). Also present is the PLK-1 activator SPAT-1/Bora (Tavernier *et al*, 2015). In contrast, just three centrosomal proteins make the list, HYLS-1, PCMD-1, and SAS-7, like PLK-1 all centriolar proteins. HYLS-1 has no known link to PLK-1 or PCM recruitment (Dammermann *et al*, 2004). However, both SAS-7 (Sugioka *et al.*, 2017) and PCMD-1 (Erpf *et al.*, 2019) are required for centrosomal PLK-1 recruitment, directly or indirectly via SPD-2. TurboID thus also identifies meaningful proximity interactors of dynamically localized proteins, although the higher number of hits overall makes it difficult to isolate novel proteins without additional information to prioritize specific candidates. We conclude that TurboID successfully identifies cytoskeletal protein interactors in *C. elegans*, albeit with notably better results for stably associated centrosomal components.

### GFP nanobody-directed TurboID as an improved method for proximity-dependent labeling

Direct TurboID tagging as described above carries with it certain disadvantages. First, to eliminate common contaminants, two TurboID strains need to be generated for each target protein, requiring a significant investment of time and effort. Moreover, those two TurboID constructs will usually be expressed as transgenes. While endogenous tagging can be carried out (Artan *et al.*, 2021), it is then difficult to generate a corresponding control matching the expression level of the TurboID-protein fusion, important for normalization of the MS data. Expression level is a particular concern when aiming to isolate proximity interactors of a protein in a specific cell type or tissue, since the relevant promoters will not necessarily match the expression level of the endogenous protein. This is assuming that the fusion protein will be able to integrate into its normal cellular context, which for centriolar proteins, often stably incorporated in previous cell cycles, will not necessarily be the case (Dammermann *et al*, 2009). Our aim was to develop an optimized tool for proximity-dependent labeling that minimizes strain generation and is able to utilize endogenous proteins, even in a cell/tissue-specific context. Rather than directly tagging the protein of interest, our approach was to express a TurboID fusion as a transgene under a ubiquitous or tissue-specific promoter and target it to the (endogenously GFP-tagged) protein of interest using a GFP nanobody (Rothbauer *et al*, 2006) (Fig. 2A, S1). Such endogenous GFP fusions will frequently already be available for proteins under study. The same GFP nanobody-TurboID promoter construct can then be used for multiple different target proteins by crossing it into the relevant GFP strain background and, when expressed alone, serves as an expression-matched control for mass spectrometry sample normalization. It should be noted that a similar strategy was recently developed for use in zebrafish (Xiong *et al*, 2021), albeit employing a conditionally stabilised GFP-binding nanobody, with implications for sample normalization as discussed below.

**Figure 2:**
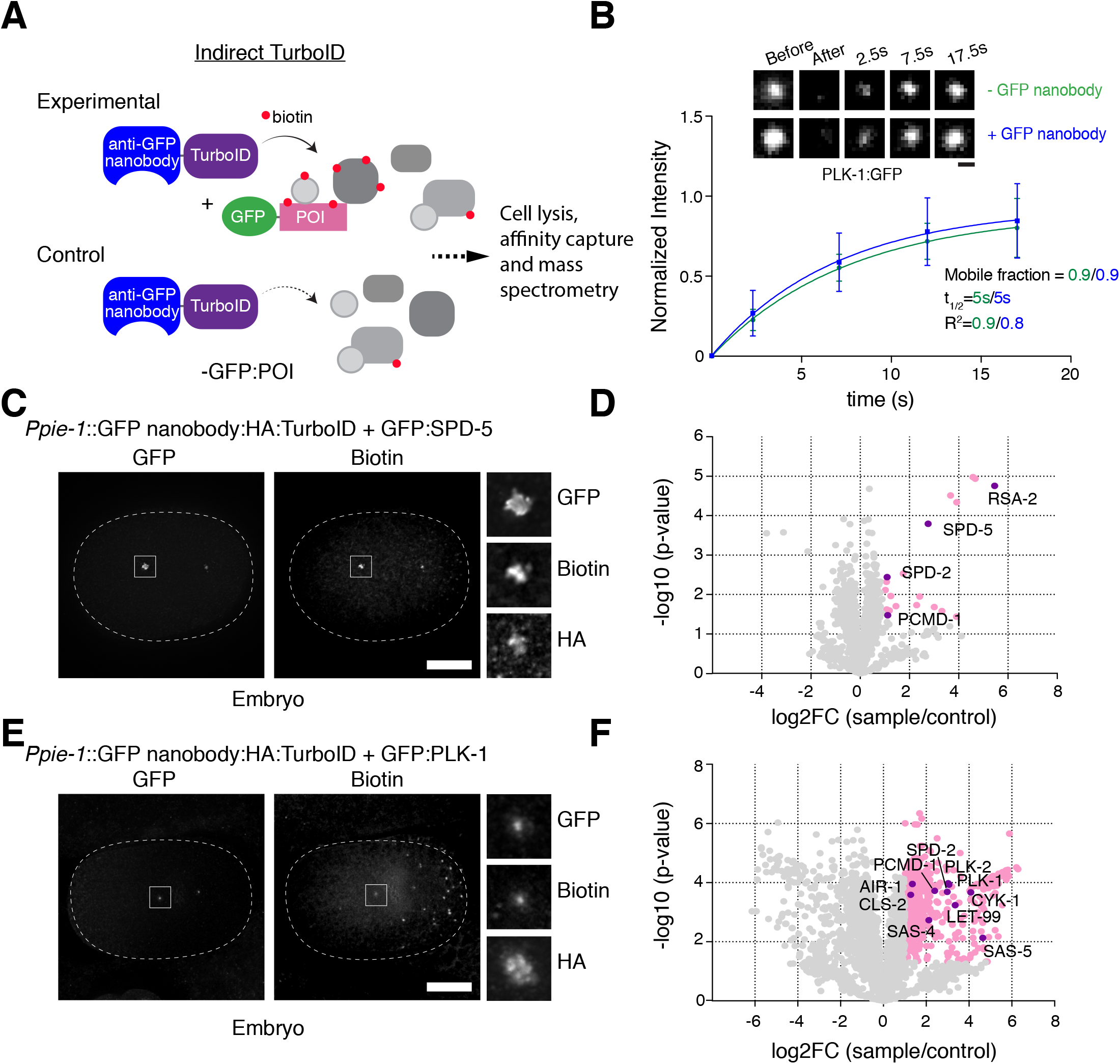
GFP nanobody-directed TurboID as an improved method for proximity-dependent labeling. (**A**) Schematic of indirect TurboID method, whereby the biotin ligase is targeted to an endogenous GFP fusion via a GFP nanobody (Rothbauer *et al.*, 2006). Note that experimental and control strains utilize the same TurboID fusion, which may be expressed under a tissue or developmental stage-specific or inducible promoter, while the target protein is potentially expressed in a wide array of tissues and cell types. (**B**) GFP nanobody addition does not perturb PLK-1 mobility. Selected images and quantitation for fluorescence recovery after photobleaching (FRAP) analysis performed on PLK-1:GFP at centrosomes in prometaphase-stage embryos in the presence (n=16 animals) or absence (n=13) of the GFP nanobody:TurboID fusion. (**C**) Indirect TurboID applied to SPD-5. Immunofluorescence micrograph of early embryo from strain co-expressing a GFP nanobody:HA:TurboID fusion under the germline promoter *pie-1* and endogenously GFP-tagged SPD-5 stained for GFP, biotin (streptavidin) and HA. Biotinylation signal is observed at centrosomes coincident with GFP:SPD-5 and the TurboID fusion. (**D**) Result of LC-MS/MS analysis for indirect TurboID on SPD-5 from mixed-stage embryos. Volcano plot of −log10 p-values against log2 fold change (sample/control). Significantly enriched proteins (Log2 enrichment >1, p-value <0.05) are indicated in pink, with selected proteins highlighted. Compare Fig. 1D. See also Supplemental Table 1 and Fig. S2E. (**E**) Indirect TurboID applied to PLK-1. Immunofluorescence micrograph of early embryo from strain co-expressing a GFP nanobody:HA:TurboID fusion under the germline promoter *pie-1* and endogenously GFP-tagged PLK-1 stained for GFP, biotin (streptavidin) and HA. Biotinylation signal is observed at centrosomes coincident with PLK-1:GFP and the TurboID fusion. (**F**) Result of LC-MS/MS analysis for indirect TurboID on PLK-1 from mixed-stage embryos. Compare Fig. 1F. See also Supplemental Table 1 and Fig. S2E. Scale bars are 1µm (B), 10µm (C, E).

To test our approach, we generated a strain expressing the GFP nanobody-TurboID fusion under the germline promoter *pie-1* and crossed it with strains expressing endogenously GFP-tagged SPD-5 and PLK-1. In both cases, immunofluorescence microscopy showed biotinylation signal colocalizing with the GFP-tagged protein, in the case of SPD-5 in the PCM and in the case of PLK-1 at centrioles, kinetochores and the spindle midzone (Fig. 2C, E). The GFP nanobody therefore successfully penetrates the centrosome and directs the biotin ligase to the appropriate intracellular location. The GFP nanobody-TurboID fusion is fairly large (a combined 50kDa for nanobody and ligase), which when added to the 27kDa of GFP could perturb the dynamics or functionality of the target protein. However, both double strains were found to be fully viable (Embryonic viability 98.6%, n=1209, GFP:SPD-5 and GFP nanobody-TurboID; 99.6%, n=1614, GFP:SPD-5 alone; 99.6%, n=2355, GFP:PLK-1 and GFP nanobody-TurboID; 98.4%, n=1512, GFP:PLK-1 alone). In addition, we assessed the dynamics of cytoplasmic exchange of GFP:PLK-1 at centrosomes by fluorescence recovery after photobleaching in prometaphase-stage embryos in the presence and absence of the nanobody. As previously reported for vertebrate somatic cells(Kishi *et al.*, 2009; Mahen *et al.*, 2011) cytoplasmic exchange of PLK-1 is extremely rapid, with a half-time of recovery t_1/2_ of 5s, with essentially no stably associated centriolar or PCM population (mobile fraction A=0.9). These dynamics remained unchanged in the presence of the nanobody (Fig. 2B). Nanobody addition therefore does not appreciably hinder localization, dynamics or functionality of the target protein.

While we were able to detect biotinylation signal at centrosomes with both GFP fusions, we sought to compare labeling efficiency of direct and indirect TurboID approaches. We therefore prepared cytoskeletal fractions from embryo extracts and performed mass spectrometry on streptavidin pulldowns as for direct TurboID, with three independent replicates for both experimental (GFP nanobody-TurboID and GFP:SPD-5/PLK-1) and control (GFP nanobody-TurboID only) conditions. One might expect that the insertion of two additional elements between biotin ligase and target protein (GFP nanobody and GFP) would widen the resultant labeling radius around the target protein. However, results were broadly similar compared to direct TurboID. In the case of SPD-5, the list of proteins significantly enriched in experimental samples over controls actually shrunk, to 22 proteins (Fig. 2D, Supplemental Table 1). With one exception (SAS-7, not detected in these mass spec samples) the same centrosomal proteins were again found to be enriched, SPD-5 itself, RSA-2, SPD-2 and PCMD-1, with the remainder of the list either uncharacterized proteins or likely contaminants. Only one of these proteins was found on both lists, B0001.2, a nematode-specific protein reported to be required for embryonic development (Piano *et al*, 2002) but otherwise currently uncharacterized. For PLK-1, the list was even longer than for direct TurboID, 501 proteins, again comprising many proteins involved in cytoskeletal organization and morphogenesis (Fig. 2F, Supplemental Table 1). Notably, only 22 proteins were common to both lists, including PCMD-1 (HYLS-1 and SAS-7 were not detected in these samples). There were, however, several additional centrosomal components, including SAS-4 (CPAP/CenpJ in vertebrates), a PLK1 target in mitotic entry (Ramani *et al*, 2018), and SPD-2 and AIR-1, involved in centriolar recruitment and activation of PLK-1 (Cabral *et al*, 2019; Joukov *et al*, 2010; Joukov *et al*, 2014). GFP nanobody-direct TurboID thus performs similarly to direct TurboID in identifying functionally relevant proximity interactors.

### Efficient tissue-specific labeling using GFP nanobody-directed TurboID

Our goal in developing GFP nanobody-direct TurboID was not only to simplify and streamline proximity labeling by utilizing available GFP fusions, but to be able to better probe protein-protein interactions in a tissue- or cell-specific context. To evaluate the suitability of our tool for these purposes, we turned to post-mitotic sensory neurons and the acentriolar centrosome at the ciliary base, whose assembly and function we recently investigated (Garbrecht *et al.*, 2021). For this we generated a strain expressing the GFP nanobody-TurboID fusion under the ciliated neuron-specific promoter *osm-6* and crossed it into the strain expressing endogenously GFP-tagged SPD-5 (PLK-1 not being expressed in post-mitotic sensory neurons (Garbrecht *et al.*, 2021)). As in the early embryo, immunofluorescence microscopy showed biotinylation signal colocalizing with the GFP-tagged protein at the ciliary base (Fig. 3A). The extended dendrite of these neurons (60µm in the L1 larval stage and nearly double that in the adult worm (Heiman & Shaham, 2009)) therefore does not present an obstacle for nanobody targeting.

**Figure 3:**
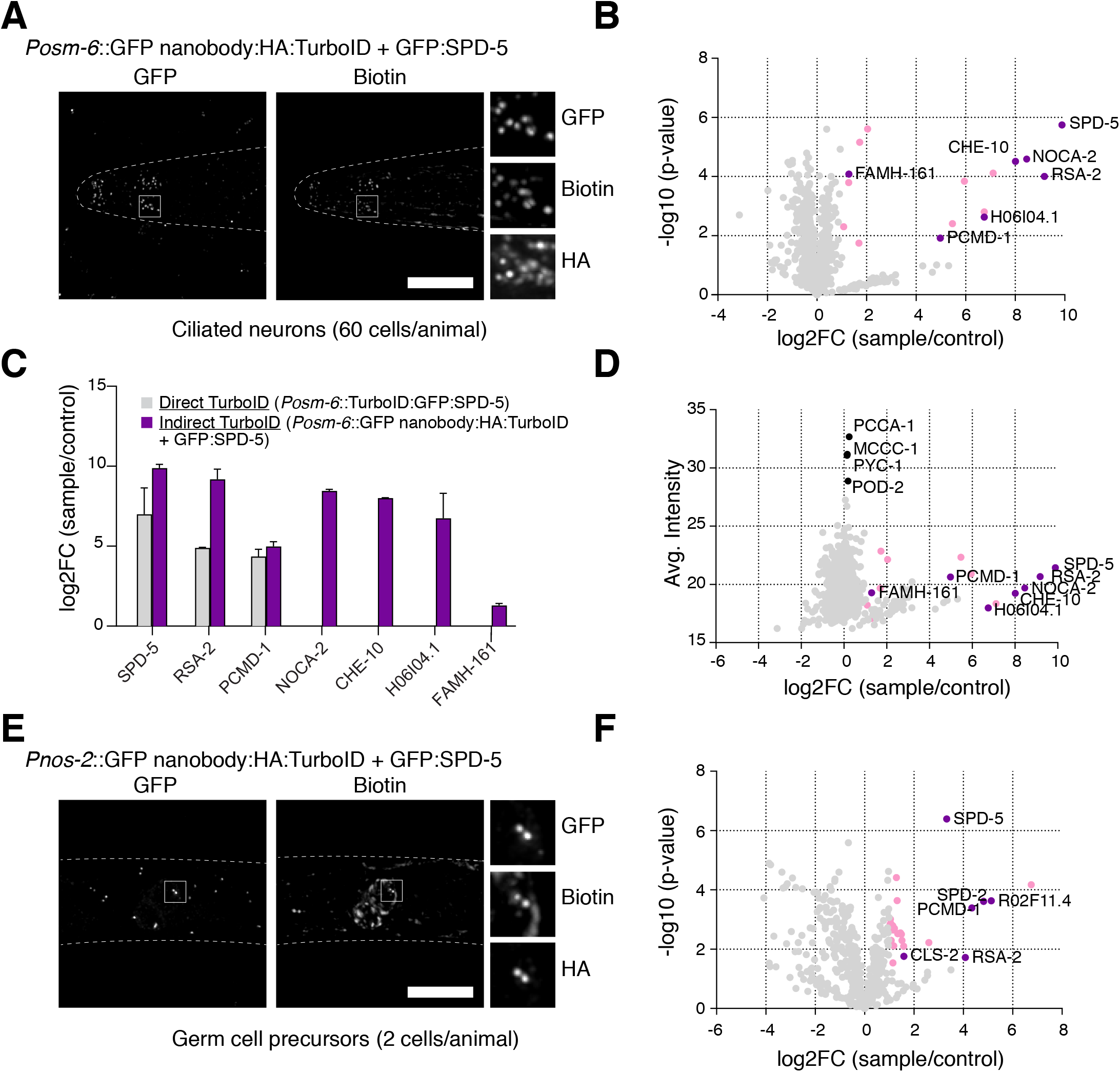
Tissue-specific labeling using GFP nanobody-directed TurboID. (**A**) TurboID applied to SPD-5 in ciliated neurons. Immunofluorescence micrograph of head of L1 larva from strain co-expressing a GFP nanobody:HA:TurboID fusion under the ciliated neuron-specific promoter *osm-6* and endogenously GFP-tagged SPD-5 stained for GFP, biotin (streptavidin) and HA. Biotinylation signal is observed at the ciliary base coincident with GFP:SPD-5 and the TurboID fusion. (**B**) Result of LC-MS/MS analysis for indirect TurboID on SPD-5 in ciliated neurons. Volcano plot of −log10 p-values against log2 fold change (sample/control). Significantly enriched proteins (Log2 enrichment >1, p-value <0.05) are indicated in pink, with selected proteins highlighted. See also Supplemental Table 1. (**C**) Comparison of LC-MS/MS results for direct and indirect TurboID on SPD-5 in ciliated neurons. Indirect TurboID identifies several additional SPD-5 proximity interactors. Full results for direct TurboID presented in Fig. S2A, B and Supplemental Table 1. (**D**) Volcano plot of log2 abundance (sample) against log2 fold change (sample/control) for indirect TurboID on SPD-5 in ciliated neurons. Significantly enriched proteins (Log2 enrichment >1, p-value <0.05) are indicated in pink, with selected proximity interactors highlighted. Note that these interactors are present at much lower levels in the sample compared to endogenously biotinylated proteins, most notably the carboxylases PCCA-1, PYC-1, POD-2 and MCCC-1 (Artan *et al.*, 2021), highlighting the importance of proper sample normalization. (**E**) TurboID applied to SPD-5 in germ cell precursors. Immunofluorescence micrograph of the area of the blast cells in an L1 larva co-expressing a GFP nanobody:HA:TurboID fusion under the germline-specific promoter *nos-2* and endogenously GFP-tagged SPD-5 stained for GFP, biotin (streptavidin) and HA. Biotinylation signal is observed at centrosomes in two cells (the primordial germ cells Z2, Z3) coincident with GFP:SPD-5 and the TurboID fusion. (**F**) Result of LC-MS/MS analysis for indirect TurboID on SPD-5 in germ cell precursors. See also Supplemental Table 1 and Fig. S2H. Scale bars in C and E are 10µm.

Ciliated neurons account for only 60 of the 959 somatic cells of the adult hermaphrodite (Inglis *et al*, 2007), not counting the two gonad arms which harbor numerous germline nuclei within a common syncytium, maturing oocytes, sperm and fertilized embryos. To maximize ciliated neuron representation within the starting material for TurboID, we therefore chose the L1 larval stage (558 cells total (Sulston & Horvitz, 1977)), a stage where ciliogenesis is largely complete (Serwas *et al*, 2017) and which can furthermore be obtained in a highly synchronized manner and in large quantities by hatching embryos in liquid culture in the absence of food (Johnson *et al*, 1984). Conditions for centrosome isolation from later developmental stages of the worm have not been established. We therefore performed TurboID on whole cell extracts as carried out by Artan et al. (Artan *et al.*, 2021). Starved worms were fed with OP50.1 supplemented with 1mM biotin for 2 hours prior to extract preparation to stimulate centrosome assembly (Garbrecht *et al.*, 2021) and enhance TurboID-dependent biotinylation (Artan *et al.*, 2021). Mass spectrometry revealed a clearly defined set of 16 proximity interactors above the significance threshold, with few obvious contaminants (Fig. 3B, Supplemental Table 1). These included RSA-2 and PCMD-1, also found as SPD-5 interactors in early embryos, but not SPD-2, a protein lost from centrosomes early in neuronal differentiation (Serwas *et al*, 2016). The remainder of the list is comprised for the most part of uncharacterized proteins known from high-throughput studies to be expressed in sensory neurons. The one exception is CHE-10, the *C. elegans* homolog of rootletin, which forms the ciliary rootlet in the region of the degenerated basal body (Mohan *et al*, 2013) in close proximity to the acentriolar centrosome. The other proteins, including FAMH-161, the apparent homolog of vertebrate FAM-161A (Zach *et al*, 2012) and B, and NOCA-2, related to vertebrate ninein-like protein/Nlp (Casenghi *et al*, 2003), must be considered strong candidates for ciliary base localization and potential basal body/ciliary centrosome function. Both FAM-161A and Nlp have been shown to play important roles in microtubule nucleation, stability and anchoring in the context of the interphase centriole/centrosome/basal body in vertebrates (Casenghi *et al.*, 2003; Di Gioia *et al*, 2012; Ganier *et al*, 2018; Le Guennec *et al*, 2020), functions that may be conserved in *C. elegans*. Another candidate protein identified by mass spectrometry is H06I04.1, a putative ortholog of CEP135/Bld10 (see below). In an effort to further compare direct and indirect GFP nanobody-directed TurboID, direct TurboID was performed in parallel using the same *osm-6* promoter to drive expression of GFP:TurboID:SPD-5 and the corresponding GFP:TurboID control. Here, direct TurboID performed markedly worse, identifying solely RSA-2 and PCMD-1 as well as the uncharacterized proteins F53G12.4, F48E3.9 and ZK809.5 (all also identified by indirect TurboID) amongst a list of 55 proximity interactors, with the remainder clear contaminants (Fig. 3C, S2A, B, Supplemental Table 1). Indirect TurboID therefore offers clear advantages over conventional direct labeling in this tissue-specific context.

### Successful tissue-specific labeling in a limited number of cells

Interaction proteomics is particularly challenging where those interactions occur in a complex source material such as within a limited number of cells in a tissue or animal. We next sought to investigate whether our TurboID was able to detect such interactions using the gonad precursor cells in L1 larvae as a test case. These are two sets of two cells, the somatic gonad precursor cells Z1 and Z4 and the primordial germ cells Z2 and Z3, which through a series of divisions at later developmental stages will give rise to the somatic gonad and germline in the adult, respectively. At the L1 larval stage these blast cells are arrested in G1 and G2 phase of the cell cycle, with what appear to be canonical interphase centrosomes (Fukuyama *et al*, 2006; Hong *et al*, 1998). With centrioles and centrosomes degenerating following terminal differentiation in most somatic tissues (Lu & Roy, 2014; Serwas *et al.*, 2017), these cells therefore represent potentially the best model for interphase centrosome assembly in the worm. As a first step towards investigating centrosome composition in those cells, we generated GFP nanobody-TurboID fusions under the control of promoters reported to be active in those cells (*Pehn-3a* Z1/Z4 (Mathies *et al*, 2019), *Pnos-2* Z2/Z3 (Lee *et al*, 2017)) and crossed them into the strain expressing GFP-tagged SPD-5. Immunofluorescence microscopy showed robust centrosomal biotinylation signal for the *Pnos-2* promoter in Z2/Z3 cells (Fig. 3E). However, for *Pehn-3a* in Z1/Z4 cells there was considerable cytoplasmic background signal, suggesting a weaker promoter would need to be used (not shown). We therefore proceeded only with *Pnos-2* in the primordial germ cells, performing TurboID on whole cell extracts prepared from L1 larvae as described above. Mass spectrometry revealed a set of 24 proximity interactors above the significance threshold. Remarkably, aside from two likely contaminants (titin, ttn-1, and mca-3, a muscle plasma membrane Ca2+ ATPase) the top hits were exclusively centrosomal proteins: SPD-2, PCMD-1, RSA-2 and CLS-2/CLASP (Espiritu *et al.*, 2012), as well as R02F11.4, a potential homolog of Cep97 (Fig. 3F, Supplemental Table 1). We conclude that GFP nanobody-directed TurboID successfully identifies protein proximity interactors, even when interactions occur only in a very limited subset of cells (in this case 2 out of 558 cells). Given the robustness of this dataset compared even with TurboID performed in embryos we propose that sample enrichment (e.g., by fractionation) is not required and may indeed result in less reliable results due to increased variability between replicates.

### Cep97 and Cep135/Bld10 as tissue-specific centrosomal components

Cep97 and Cep135/Bld10 are well-known centriolar components with critical roles in centriole assembly/stability and centrosome/cilium biogenesis (Carvalho-Santos *et al*, 2012; Dobbelaere *et al*, 2008; Dobbelaere *et al*, 2020; Fu *et al*, 2016; Matsuura *et al*, 2004). Cep135 in particular is highly conserved and along with SAS-4 and SAS-6 has been proposed to be one of three proteins defining centriole/basal body architecture across eukaryotes (Carvalho-Santos *et al*, 2010). It is then surprising that both Cep97 and Cep135 have been reported to be absent from the *C. elegans* genome (Carvalho-Santos *et al.*, 2010; Hodges *et al*, 2010). However, it must be noted that nematode genomes are highly divergent such that even conserved components such as PLK4/ZYG-1 were originally reported to be missing based on lack of amino acid sequence homology (Carvalho-Santos *et al.*, 2010; O’Connell *et al*, 2001) and not recognized as such until much later based on conserved 3D structure and molecular function. Their identification has thus been almost invariably based on genome-wide screens performed for centriole/centrosome assembly in the early embryo and somewhat less comprehensive genetic screens for cilium biogenesis in sensory neurons (Inglis *et al.*, 2007; Oegema & Hyman, 2006).

The identification of R02F11.4 as the ortholog of Cep97 is quite clear to the extent of being annotated as such on Wormbase (Version WS282, November 2021), although it does require the use of intermediary, less divergent, species for successive reciprocal BLAST identification or Position-Specific Iterated BLAST (PSI-BLAST) analysis (Fig. S3). 3-dimensional structure prediction via AlphaFold (Jumper *et al*, 2021) also suggests a similar protein architecture of the *C. elegans*, *Drosophila* and human proteins, including a leucine rich repeat region and extended coiled coils (Fig. 4A). Generation of an endogenous promoter GFP transgene revealed CEP-97 localization to centrioles in late-stage embryos, gonad precursors and the ciliary base, though curiously not early embryos (Fig. 4B). This is consistent with the lack of a reported early embryo RNAi phenotype in high-throughput screens, a result we have confirmed in our own hands (Embryonic viability 99.6%, n=1219, *R02F11.4(RNAi)*; 1.1%, n=1272, *spd-5(RNAi)*; 99.8%, n=1238, Control). However, a partial gene deletion results in fully penetrant L1 larval arrest, suggesting a critical role in later development. Further experiments will be necessary to define CEP-97 function in the worm. However, it is striking to note that CEP-97 is first detectable at centrioles as the cell cycle slows in late-stage embryos and strongly accumulates at centrioles in gonad precursor cells, cells which as previously noted are arrested for extended periods of time in interphase. These findings are consistent with recent reports of cell cycle-entrained oscillations of Cep97 levels regulating centriole growth in *Drosophila* (Aydogan *et al*, 2021).

**Figure 4:**
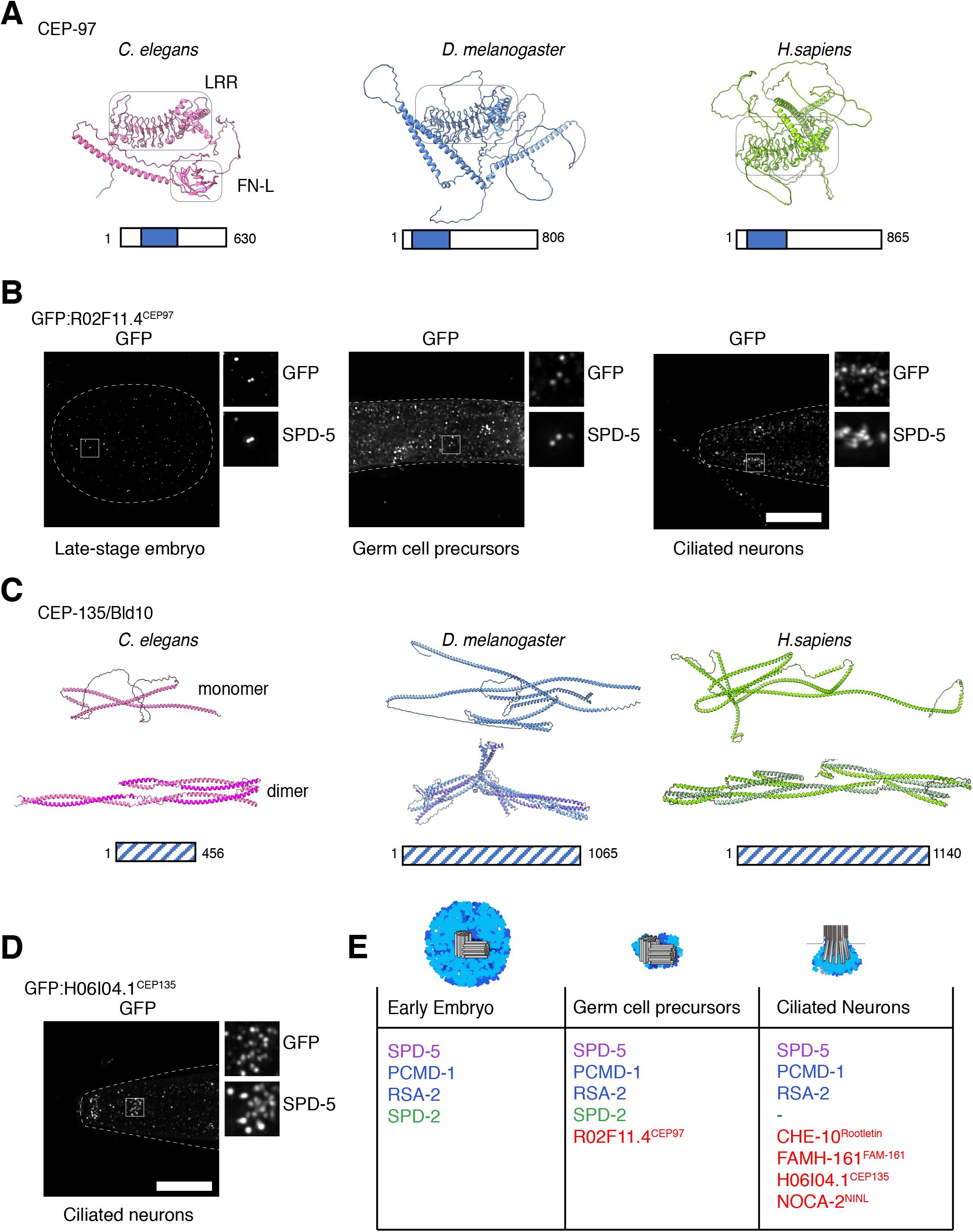
Cep97 and Cep135/Bld10 as novel tissue-specific centrosomal components in *C. elegans.* (**A**) AlphaFold (Jumper *et al.*, 2021) predicts a similar 3D architecture for *C. elegans* R02F11.4 and its putative *Drosophila* and human orthologs Cep97, including a prominent leucine rich repeat that is highly conserved across Cep97 orthologs followed by long coiled coils (see Fig. S3). Additionally, a Fibronectin type III (FN-3) domain is predicted for the *C. elegans* protein, which is conserved in other nematode homologs but not in insects or vertebrates. (**B**) An endogenous promoter GFP transgene shows R02F11.4 expressed in late-stage embryos, germ cell precursors and ciliated neurons, localizing to centrioles (marked by immunofluorescence co-staining with SPD-5) and the ciliary base (likewise), respectively. (**C**) AlphaFold structure prediction shows *C. elegans* H06I04.1 and its putative *Drosophila* and human orthologs Cep135/Bld10 to display a similar 3D architecture, composed primarily of coiled coils. These align in a rope-like manner when predicting the proteins to form dimers as has been shown for human and *Chlamydomonas* Cep135/Bld10 (Kraatz *et al.*, 2016). For hsCep135 and dmCEP135 only the N-terminal regions (1-600, hsCEP135; 1-547, dmCEP135) were used for homodimer prediction to reduce computational costs. We note that in the homidimeric dmCEP135 model some regions are overlapping suggesting that the native structure deviates from the prediction. For H06I04.1 the first 131 residues, predicted with lower confidence, are not displayed. See also Fig. S4. (**D**) An endogenous promoter GFP transgene shows H06I04.1 expressed exclusively in ciliated sensory neurons of the head (amphids and cephalic/labial neurons) and tail (phasmids, not shown), localizing to the ciliary base (marked by immunofluorescence co-staining with SPD-5). (**E**) Comparison of selected SPD-5 proximity interactors identified in embryos, germ cell precursors and ciliated neurons (for full list see Supplemental Table1). Common to all three tissue contexts are PCMD-1 and RSA-2, while SPD-2 is absent from post-mitotic sensory neurons. The previously uncharacterized proteins R02F11.4 and H06I04.1 are amongst the tissue-specific SPD-5 interactors, identified in germ cell precursors and ciliated neurons, respectively. Scale bars in B and D are 10µm.

H06I04.1 initially caught our attention as a putative interactor of the centriolar proteins SAS-6, FAMH-161/FAM161A/B and NOCA-2/NINL as well as the microtubule-associated protein ZYG-8/Doublecortin in high-throughput screens (Li *et al*, 2004; Simonis *et al*, 2009). Direct BLAST searches initially failed to identify any homologs outside of nematodes. However, repeated rounds of Position-Specific Iterated BLAST (PSI-BLAST) identified homologs first in insects and then vertebrates, all related to Cep135/Bld10 (Fig. S4). Amino acid sequence conservation is clearly very low. However, *C. elegans*, *Drosophila* and human proteins all contain multiple predicted coiled coils, which for human Cep135 have been shown to be important for dimerization and microtubule binding (Kraatz *et al*, 2016) (Fig. 4C). An endogenous promoter GFP transgene revealed CEP-135 expression exclusively in post-mitotic neurons, where it localizes to the ciliary base as well as the nerve ring (Fig. 4D and not shown). No clear signal was observed elsewhere in the worm, consistent with the lack of a reported RNAi phenotype in high-throughput screens and our own hands (Embryonic viability 99.9%, n=1715, *H06I04.1(RNAi)*). CEP-135 would thus appear to function specifically in the context of centriole-derived basal bodies. While Cep135 is generally thought of as a cartwheel component important for centriole assembly (Matsuura *et al.*, 2004), a more limited function in the context of cilia would be consistent with work in *Drosophila*, where it is basal bodies and consequently cilia that are primarily affected in Cep135 mutants (Bayless *et al*, 2012; Carvalho-Santos *et al.*, 2012; Matsuura *et al.*, 2004).

### Conclusions and outlook

Proximity labeling by BioID has provided valuable insights into the molecular composition of centrosomes and related structures in vertebrate cells (Firat-Karalar *et al.*, 2014; Gheiratmand *et al.*, 2019; Gupta *et al.*, 2015). The recent development of TurboID (Branon *et al.*, 2018) promises to do the same for invertebrate experimental models like *C. elegans* and *Drosophila*. The indirect GFP nanobody-directed TurboID approach developed here represents a refinement of this method, which in genetic models greatly simplifies strain generation by allowing the repurposing of existing GFP-tagged strains for TurboID. More importantly, when not combined with the GFP fusion of interest the GFP nanobody-TurboID strain represents the ideal expression-matched control to remove non-specific background and identify *bona fide* proximity interactors in the often extensive mass spectrometry lists. This superior ability to filter mass spectrometry data is demonstrated by strikingly successful tissue-specific labeling, even where the bait is expressed in only a highly limited number of cells, such as the germ cell precursors. Abundant endogenously biotinylated proteins, including the carboxylases PCCA-1, PYC-1, POD-2 and MCCC-1 noted by Artan et al. (Artan *et al.*, 2021), do represent a significant proportion of the isolated peptides (see Fig. 3D), but can easily be filtered out. Enriching for a specific cell type or intracellular structure of interest as we did by fractionation of embryo extracts is therefore clearly not necessary and may actually result in less reliable results due to increased variability between replicates. As previously noted, a similar approach employing a conditionally stabilised GFP-binding nanobody was recently developed for use in zebrafish (Xiong *et al.*, 2021). In this approach, the GFP nanobody-TurboID fusion is destabilized when not bound to the protein of interest, reducing non-specific labeling. However, this also compromises the ability to use the GFP nanobody-TurboID fusion alone as an expression-matched control for mass spectrometry data normalization. Regardless of their individual advantages and disadvantages, both strategies clearly hold great promise in tackling previously inaccessible problems in interaction proteomics across genetic models from *C. elegans* to vertebrates. The use of inducible promoters in combination with indirect TurboID should further expand the scope of possible proximity labeling experiments, something we have not explored here.

Contrary to common perception, proximity labeling by TurboID appears to be highly restricted, identifying only those proteins in closest proximity to the bait protein. In the case of SPD-5, these are proteins like PCMD-1, SPD-2 and RSA-2, all of which are known to interact with SPD-5 and regulate PCM assembly (see below). Other PCM proteins including highly abundant ones like γ-tubulin are not detected, suggesting a labeling radius even more narrow than the 10-15 nm previously estimated for BioID (Kim *et al.*, 2014). This method is therefore ideally suited for the identification of potential protein-protein interactors, but perhaps less so for compartment labeling, even when using a bait protein found throughout a structure of interest such as the PCM. In comparison with our results for SPD-5, a stably bound component of the PCM scaffold (Laos *et al.*, 2015), our results for the dynamically localized kinase PLK-1 were markedly less clear, with extensive background requiring strategies to prioritize hits and identify true proximity interactors. Performing additional replicates or combining multiple baits may help make sense of such more complex data.

In this proof of principle study, we applied our tool to carry out an initial characterization of tissue-dependent variation in PCM composition in the worm, focusing on proximity interactors of the scaffolding component SPD-5 in embryos, germ cell precursors and at the ciliary base (Fig. 4E). Our data identify RSA-2 and PCMD-1 as SPD-5 proximity interactors common to all three developmental contexts. PCMD-1 is perhaps not a surprise, having been found to contribute to SPD-5 recruitment/tethering to centrioles in the early embryo and the ciliary base in sensory neurons (Erpf *et al.*, 2019; Garbrecht *et al.*, 2021; Stenzel *et al*, 2021), RSA-2 somewhat more so. While RSA-2 was originally identified as a centrosomal PP2A recruitment factor (Schlaitz *et al.*, 2007), its proximity interaction with SPD-5 at all types of centrosomes in the apparent absence of the phosphatase suggests a more direct role in the regulation of PCM assembly, which bears closer investigation. There is also considerable tissue-dependent variation in PCM composition. Thus, SPD-2 is only detected in mass spectrometry samples for SPD-5 in embryos and germ cell precursors, but not sensory neurons, consistent with the previously reported loss of SPD-2 from the acentriolar centrosome at the ciliary base (Garbrecht *et al.*, 2021; Serwas *et al.*, 2017). Other proteins are only detected at later developmental stages, such as Cep97. Its appearance at centrioles in late-stage embryos may reflect a slowing of the cell cycle allowing its accumulation during the more extended interphase (Aydogan *et al.*, 2021), and be connected to the emergence of centriolar doublet microtubules at approximately the same time (Nechipurenko *et al*, 2017; Serwas *et al.*, 2017). Finally, ciliated sensory neurons present a unique set of SPD-5 proximity interactors, including CHE-10/rootletin. A surprising finding here is the tentative identification of a homolog of Cep135/Bld10, previously thought to be missing in nematodes, present exclusively at the ciliary base. Further work will be needed to clarify the functional contribution of this protein to basal body architecture and cilium assembly in *C. elegans*, but this tissue-specific expression pattern is consistent with a role for this protein specifically in the context of cilia (Bayless *et al.*, 2012; Carvalho-Santos *et al.*, 2012; Matsuura *et al.*, 2004).

In summary, then, our data suggest that centrioles and centrosomes may be much more variable across different tissues and developmental stages than currently appreciated. The systematic application of newly developed tools such as tissue-specific TurboID and inducible degron-mediated degradation to investigate their molecular composition and the underlying molecular mechanisms at work should help shed light on what has hitherto been a somewhat neglected aspect of centrosome biology.

## MATERIALS AND METHODS

### EXPERIMENTAL MODEL AND SUBJECT DETAILS

#### *C. elegans* strains and culture conditions

*C. elegans* strains expressing endogenously tagged GFP:SPD-5 (Cabral *et al.*, 2019) and PLK-1:GFP (Martino *et al*, 2017) have been described previously. Strains expressing endogenous promoter-driven GFP:TurboID:SPD-5 and GFP:TurboID:PLK-1 and *Posm-6*-driven TurboID:GFP:SPD-5, as well as corresponding control strains lacking the SPD-5/PLK-1 coding sequence, were generated by cloning the corresponding genomic locus including 5’ and 3’ regulatory sequences, GFP and TurboID into the targeting vector pCFJ151, followed by Mos1-mediated transposition (Frokjaer-Jensen *et al.*, 2008). SPD-5 and GFP coding sequence were derived from plasmid pAD395 (Cabral *et al*, 2013), TurboID from plasmid pAS31 (Branon *et al.*, 2018). All other fragments were amplified from *C. elegans* genomic DNA (length of *spd-5* promoter 1150bp, 3’UTR 577bp; *plk-1* promoter 2000bp, 3’UTR 2324bp; *osm-6* promoter 425bp). Strains expressing the GFP nanobody-TurboID fusion under various tissue-specific 5’ and 3’ regulatory sequences were generated by Mos1 transposon insertion as above. HA tagged-TurboID was derived from plasmid pAS31(Branon *et al.*, 2018), the GFP nanobody from plasmid pAD834(Garbrecht *et al.*, 2021). Promoters and 3’UTRs were amplified from plasmids pCFJ152 (*pie-1* promoter, 1105bp, (Frokjaer-Jensen *et al.*, 2008)) and pAD834 (*tbb-2* 3’UTR, 330bp (Garbrecht *et al.*, 2021)) as well as *C. elegans* genomic DNA (*osm-6* promoter 425bp; *nos-2* promoter, 956bp, 3’UTR, 1000bp; *ehn-3a* promoter, 2004bp, 3’UTR 806bp). *pie-1* promoter-driven GFP nanobody-TurboID is also combined with mCherry:histone H2B as a visible marker co-expressed via an operon linker, amplified from plasmid pLC754 (gift from Luisa Cochella, (Charest *et al*, 2020)). Strains expressing endogenous promoter-driven GFP:CEP-97 and GFP:CEP-135 were generated by Mos1 transposon insertion as above. In each case promoter (*cep-97* 609bp, *cep-135* 1984bp) and 3’UTR (*cep-97* 491bp, *cep-135* 910bp) were amplified from *C. elegans* genomic DNA, coding sequence from cDNA. All constructs described above were assembled by Gibson assembly (Gibson *et al*, 2009). Strains co-expressing endogenously GFP-tagged SPD-5/PLK-1 and GFP nanobody-TurboID expressed under the different tissue-specific promoters were constructed by mating. The genotypes of all strains used are listed in the Reagents and Tools Table. All strains were maintained at 23°C.

### METHOD DETAILS

#### Sequence analysis and 3D structure predictions

Vertebrate orthologs of uncharacterized proteins in our datasets were identified by reciprocal BLAST analysis using the *C. elegans* protein as the starting point. Where direct comparisons failed to identify a clear homolog, indirect searches were performed using less divergent related species as intermediates. Multiple sequence alignments were generated using MUSCLE within Jalview (http://www.jalview.org), while phylogenetic trees were constructed also in Jalview by neighbor joining using the BLOSUM62 matrix. 3D structure predictions of monomeric proteins were obtained from the AlphaFold database (Jumper *et al.*, 2021). Prediction of homodimeric assemblies of CEP135 homologs was performed with AlphaFold-multimer (Evans *et al*, 2021). Molecular graphics of the structures were created using UCSF ChimeraX (Goddard *et al*, 2018).

#### Live Imaging

Live imaging of embryos and L1 larvae was performed on a Zeiss Axio Imager Z2 microscope equipped with a 63x 1.4NA Plan Apochromat objective and Lumencor SOLA SE II light source. Single plane GFP fluorescence and DIC images were acquired using a Photometrics CoolSNAP-HQ2 cooled CCD camera controlled by ZEN 2 blue software (Zeiss) and imported into Fiji for post-acquisition processing.

#### Immunofluorescence and Fixed Imaging

Immunofluorescence experiments were performed as previously described (Oegema *et al*, 2001) using affinity-purified antibodies to GFP, SPD-5, TAC-1 and HA, as well as flourorescently labeled streptavidin to detect biotinylated proteins. *C. elegans* embryos and L1 larvae were permeabilized by freeze-crack, fixed in −20°C methanol for 20min, rehydrated in PBS, blocked in AbDil (PBS, 2% BSA, 0.1% Triton X-100) for 20min, incubated with 5µg/ml Streptavidin-Alexa Fluor 647 and directly labeled or unlabeled primary antibodies at 1μg/ml in AbDil for 2h (for mouse anti-HA.11 overnight at 4°C), washed 3x in PBST, incubated with secondary antibodies at 7.5μg/ml in AbDil for 1h followed by Hoechst 33342 at 1µg/ml in PBS for 5min, washed in PBST and mounted in Vectashield. 3D widefield datasets were acquired using a 100x 1.4NA Uplan S Apochromat objective on a DeltaVision 2 Ultra microscope equipped with 7-Color SSI module and sCMOS camera and controlled by Acquire Ultra acquisition software (GE Healthcare), computationally deconvolved using the enhanced ratio constrained iterative deconvolution algorithm, and maximum-intensity projected before being imported into Photoshop (Adobe) for panel preparation. No nonlinear gamma corrections were performed during image processing.

#### Fluorescence Recovery after Photobleaching

To examine the dynamics of PLK-1 at centrosomes in early embryos, adult worms were dissected in meiosis medium (60% Leibowitz L-15 media, 25mM Hepes pH7.4, 0.5% Inulin, 20% heat-inactivated fetal bovine serum) and embryos filmed without compression (Monen *et al*, 2005) on a Yokogawa CSU X1 spinning disk confocal mounted on a Zeiss Axio Observer Z1 inverted microscope equipped with a 63x 1.4NA Plan Apochromat objective, 120mW 405nm and 100mW 488nm solid-state lasers, 2D-VisiFRAP Galvo FRAP module and Photometrics CoolSNAP-HQ2 cooled CCD camera and controlled by VisiView software (Visitron Systems). Low laser illumination (max power of 0.32mW) was used to minimize photobleaching. Embryos in the first mitotic division were followed from early prophase, with 6×0.5µm z-series acquired without binning at irregular intervals. Photobleaching was performed in prometaphase using the galvanometer point scanner to target a region encompassing the centrosome with the 405nm laser at 10mW power and embryos imaged until completion of cytokinesis. Embryos were analyzed only if centrosome signal was completely eliminated throughout the entire z-volume.

#### TurboID-based enzymatic protein labeling and extraction of biotinylated proteins from *C. elegans* embryos

Gravid adult *C. elegans* cultivated in a 500ml liquid culture (Zanin *et al*, 2011) were bleached to harvest embryos. Embryos were washed 3x with cold M9 followed by 1 wash with lysis buffer H100 (50mM HEPES pH7.4, 1mM EGTA, 1mM MgCl2, 100mM KCl, 10% Glycerol, 0.05% NP-40) before pelleting at 800g for 2min at 4°C. Embryos were then resuspended with lysis buffer supplemented with Roche cOmplete™ Mini EDTA-free Protease-Inhibitor-Cocktail (1 tablet/10ml lysis buffer), 1mM PMSF, 1mM Benzamidine at a ratio of 1:4 packed embryos to buffer and split into three replicate samples. Embryos in each sample were then lysed by tip sonication (30% continuous output, Bandelin Sonopuls GM70) with three pulses for 15s with brief cooling on ice between pulses. Lysates were clarified by two brief spins at 200g for 3min at 4°C in a benchtop centrifuge before pelleting insoluble cellular material including centrosomes by centrifugation at 20000g for 30min. Pellets were resuspended by boiling in 60µl 2% SDS in PBS containing 1% beta-Mercaptoethanol for 30min with multiple vortexing steps in between. 9 volumes of PBS and 1 volume 20% Triton X-100 in PBS were then added and samples sonicated three times for 30s (60% output at 1 pulse/s). Another 9 volumes of PBS were then added and samples sonicated for another 30s as before centrifuging at 20000g for 30min at 4°C and recovering the supernatant. Pierce^TM^ Streptavidin-coated magnetic beads (150µl bead slurry per sample) were equilibrated with PBS containing 0.1% SDS before incubating with solubilized cytoskeletal fractions on a rotator overnight at 4°C. Unbound lysate was then removed and beads washed for 8min each on a rotator at room temperature, first with 2% SDS in ddH_2_O, then 0.1% deoxycholic acid, 1% Triton X-100, 1mM EDTA, 500mM NaCl and 50mM HEPES pH7.5, and finally 0.5% deoxycholic acid, 0.5% NP-40, 1mM EDTA, 500mM LiCl and 10mM Tris pH7.5. Beads were then washed 5x for 3min with 50mM Tris pH7.4 before being sent for on-bead protein digestion and mass spectrometry analysis.

#### TurboID-based enzymatic protein labeling and extraction of biotinylated proteins from *C. elegans* L1 larvae

Gravid adult *C. elegans* cultivated in a 500ml liquid culture (Zanin *et al.*, 2011) were bleached to harvest embryos and those embryos allowed to hatch overnight by shaking in M9 buffer without food at 22°C to obtain a synchronous culture arrested at L1 stage. For the last 2h *E. coli* OP50.1 (800µl bacterial slurry/50ml M9) and 1mM biotin were added to stimulate centrosome assembly and biotin incorporation (Artan *et al.*, 2021). L1 larvae were then washed three times with ice-cold M9 and allowed to settle on ice after the last wash in order to aspirate off the supernatant. Two volumes of RIPA buffer (1% Triton X-100, 1mM EDTA, 0.5% sodium deoxycholate, 0.1% SDS, 150mM NaCl, 50mM Tris-HCl pH7.4) supplemented with 1mM PMSF and Roche cOmplete™ Mini EDTA-free Protease-Inhibitor-Cocktail (1 tablet/25ml lysis buffer) were added to one volume of packed worms. L1 larvae were again allowed to settle on ice and added dropwise to liquid nitrogen to obtain frozen worm ‘popcorn’, which were ground to a fine powder using a SPEX 6875 cryogenic mill (Settings: 5 cycles, 1min pre-cool, 2min runtime, 1min cool-time; 12 cps) and stored at −80°C. Worm powder was thawed in a 50ml conical tube on a tube roller at 4°C and the sample collected at the bottom of the tube by a brief spin at 200g for 1min at 4°C. SDS and DTT were then added to a final concentration of 1% and 10mM, respectively. Tubes were gently inverted several times and the sample split into three 2ml microcentrifuge tubes which were immediately incubated at 90°C for 5min. After boiling, samples were sonicated by tip sonication (20% continuous output, Bandelin Sonopuls GM70) with 2 pulses of 1min with brief cooling between pulses. Samples were then adjusted to 2M urea using a stock solution of 8M urea, 1% SDS, 50mM Tris-HCl pH7.4, 150mM NaCl) and centrifuged at 100000g for 45min at 22°C. The clear supernatant between pellet and surface lipid layer was transferred to a new tube. Zeba^TM^ spin desalting columns (7K MWCO) (Thermofisher) were equilibrated three times with 5ml RIPA buffer containing 1% SDS, 2M urea and protease inhibitors as above by centrifugation at 1000g for 5min. In order to remove free biotin, clarified samples were loaded twice onto equilibrated columns and desalted by centrifugation (1000g, 5min) to remove free biotin.

Pierce^TM^ Streptavidin magnetic beads (150µl bead slurry per sample) were equilibrated with 50mM HEPES-NaOH pH7.8 containing 0.2% Tween-20 by washing them three times with the buffer. A mixture of 190µl 50mM HEPES-NaOH pH7.8 containing 0.2% Tween-20 and 10µl 100mM Pierce^TM^ Sulfo-NHS-Acetate (dissolved in DMSO) was used to acetylate free amines on streptavidin beads by incubation at room temperature for 1h. Beads were then washed three times with 50mM NH_4_HCO_3_ containing 0.2% Tween-20 before adding the desalted and clarified sample solution and incubating on a rotator overnight at room temperature. Following incubation, unbound lysate was removed and beads washed twice with 150mM NaCl, 1mM EDTA, 2% SDS, 50mM Tris-HCl, pH7.4, once with 1X TBS, twice with 1M KCl, 1mM EDTA, 50mM Tris-HCl, 0.1% Tween-20, pH7.4, twice with 0.1M Na_2_CO_3_, 0.1% Tween-20, pH11.5, twice with 2M urea, 10mM Tris-HCl, 0.1% Tween-20, pH8.0 and finally five times with 1X TBS. Beads were then resuspended in 1X TBS buffer and sent for on-bead protein digestion and mass spectrometry analysis.

#### Sample preparation for mass spectrometry analysis

Beads were resuspended in 50µl 1M urea and 50mM ammonium bicarbonate. Disulfide bonds were reduced with 2µl of 250mM dithiothreitol (DTT) for 30min at room temperature before adding 2µl of 500mM iodoacetamide and incubating for 30min at room temperature in the dark. Remaining iodoacetamide was quenched with 1µl of 250mM DTT for 10min. Proteins were digested with 300ng trypsin (Trypsin Gold, Promega) in 3µl 50mM ammonium bicarbonate followed by incubation at 37°C overnight. The supernatant without beads was transferred to a new tube, the digest stopped by addition of trifluoroacetic acid (TFA) to a final concentration of 0.5% and the peptides desalted using C18 Stagetips (Rappsilber *et al*, 2007).

#### Liquid chromatography separation coupled to mass spectrometry

Peptides were separated on an Ultimate 3000 RSLC nano-flow chromatography system (Thermo-Fisher), using a pre-column for sample loading (Acclaim PepMap C18, 2cm × 0.1mm, 5μm, Thermo-Fisher), and a C18 analytical column (Acclaim PepMap C18, 50cm × 0.75mm, 2μm, Thermo-Fisher), applying a segmented linear gradient from 2% to 35% and finally 80% solvent B (80% acetonitrile, 0.1% formic acid; solvent A 0.1% formic acid) at a flow rate of 230nL/min over 120min. The peptides eluted from the nano-LC were analyzed by mass spectrometry as described below.

For SPD-5 embryo samples, a Q Exactive HF-X Orbitrap mass spectrometer (Thermo Fisher) coupled to the column with a nano-spray ion-source using coated emitter tips (PepSep, MSWil), was used with the following settings: The mass spectrometer was operated in data-dependent acquisition mode (DDA), survey scans were obtained in a mass range of 375-1500m/z with lock mass activated, at a resolution of 120k at 200m/z and an AGC target value of 3E6. The 12 most intense ions were selected with an isolation width of 1.4m/z, fragmented in the HCD cell at 28% collision energy and the spectra recorded for max. 200ms at a target value of 1E5 and a resolution of 30k. Peptides with a charge of +2 to +6 were included for fragmentation, the peptide match and the exclude isotopes features enabled, and selected precursors were dynamically excluded from repeated sampling for 30s. Raw data were directly used for further analysis.

For all other samples, an Exploris 480 Orbitrap mass spectrometer (Thermo Fisher) coupled to the column with a FAIMS pro ion-source (Thermo-Fisher) using coated emitter tips (PepSep, MSWil), was used with the following settings: The mass spectrometer was operated in DDA mode with two FAIMS compensation voltages (CV) set to −45 or −60 and 1.5s cycle time per CV. The survey scans were obtained in a mass range of 350-1200m/z, at a resolution of 60k at 200m/z and a normalized AGC target at 100%. The most intense ions were selected with an isolation width of 1.0m/z, fragmented in the HCD cell at 28% collision energy and the spectra recorded for max. 100ms at a normalized AGC target of 100% and a resolution of 15k. Peptides with a charge of +2 to +6 were included for fragmentation, the peptide match feature was set to preferred, the exclude isotope feature was enabled, and selected precursors were dynamically excluded from repeated sampling for 45s. Raw data were split for each CV and converted to mzxml-format using freely available software (https://github.com/coongroup/FAIMS-MzXML-Generator) for further analysis.

### QUANTIFICATION AND STATISTICAL ANALYSIS

#### Image analysis

Quantification of centrosomal PLK-1 signal was performed in Fiji. GFP signal was measured on single planes at each time point. Two variable size concentric regions were drawn around the centrosome, a smaller one encompassing the centrosome and a larger one including the surrounding cytoplasm as background. The integrated GFP intensity was then calculated by subtracting the mean fluorescence intensity in the area between the two boxes (mean background) from the mean intensity in the smaller box and multiplying by the area of the smaller box. Measurements taken after photobleaching were normalized to the mean intensity of measurements shortly before photobleaching. GraphPad Prism was then used to fit the data to the exponential equation Y = A*(1-exp(-k*X))+B where A is the mobile fraction, B is the background left after the bleach, and the half life t_1/2_ = ln(0.5)/-k. R values are the correlation coefficient obtained by fitting experimental data to the model. Data points on the graphs are the mean of the normalized GFP intensity measurements.

#### Mass spectrometry database search and analysis

MS data were analyzed using the MaxQuant software package (version 1.6.17.0, (Tyanova *et al*, 2016)) and the Uniprot *Caenorhabditis elegans* reference proteome (www.uniprot.org), as well as a database of most common contaminants. The search was performed with full trypsin specificity and a maximum of two missed cleavages at a protein and peptide spectrum match false discovery rate of 1%. Carbamidomethylation of cysteine residues were set as fixed, oxidation of methionine and N-terminal acetylation as variable modifications. For label-free quantification the “match between runs” feature and the LFQ function were activated - all other parameters were left at default. MaxQuant output tables were further processed in R (R Core Team, 2018) using a script developed in-house (https://github.com/moritzmadern/Cassiopeia_LFQ). Reverse database identifications, contaminant proteins, protein groups identified only by a modified peptide, protein groups with less than two quantitative values in one experimental group, and protein groups with less than 2 razor peptides were removed for further analysis.

The analysis of differential abundance in mass spectrometry-based proteomics data sets requires the application of missing value imputation where a particular protein of interest is missing in one or more replicates (Webb-Robertson *et al*, 2015). For the data presented in the figures we decided to perform deterministic lowest of detection (LOD) within Microsoft Excel for the comparison of different runs of mass spectrometry. LOD uses the lowest measured LFQ value within a run to fill in missing LFQ values after filtering out contaminants and proteins with less than two razor and unique peptides (Gardner & Freitas, 2021). A common alternative approach presented in the Supplemental tables is random drawing from a left-censored normal distribution (ND) in logarithmic space, which we performed in R. For this, missing values were replaced by randomly drawing data points from a normal distribution modeled on the whole dataset (data mean shifted by −1.8 standard deviations, width of distribution of 0.3 standard deviations). Differences between groups were statistically evaluated using the LIMMA package (Ritchie *et al*, 2015) at 5% FDR (Benjamini-Hochberg).

While the latter approach is often appropriate when comparing eg two treatment conditions, in our case we noted that centrosomal proteins frequently were absent in one or all control samples but also not highly abundant in the experimental samples, likely because of their restricted spatial distribution and low abundance within the cell. Application of ND imputation consequently led to misleading variations in protein abundance with less clear separation of likely proximity interactors from apparent contaminants, while also frequently inverting sample enrichment ratios between hits. In contrast, we found LOD resulted in a more accurate representation of sample enrichment and this approach was used in our data presentations. In essentially all cases, however, likely proximity interactors passed the significance threshold with both methods of imputation. These were defined as a log2 fold change of >1 and a p-value in an unpaired t-tests of <0.05. GraphPad Prism was used to prepare bar graphs, volcano and MA plots using LFQ values imported from Microsoft Excel.

### REAGENTS AND TOOLS TABLE

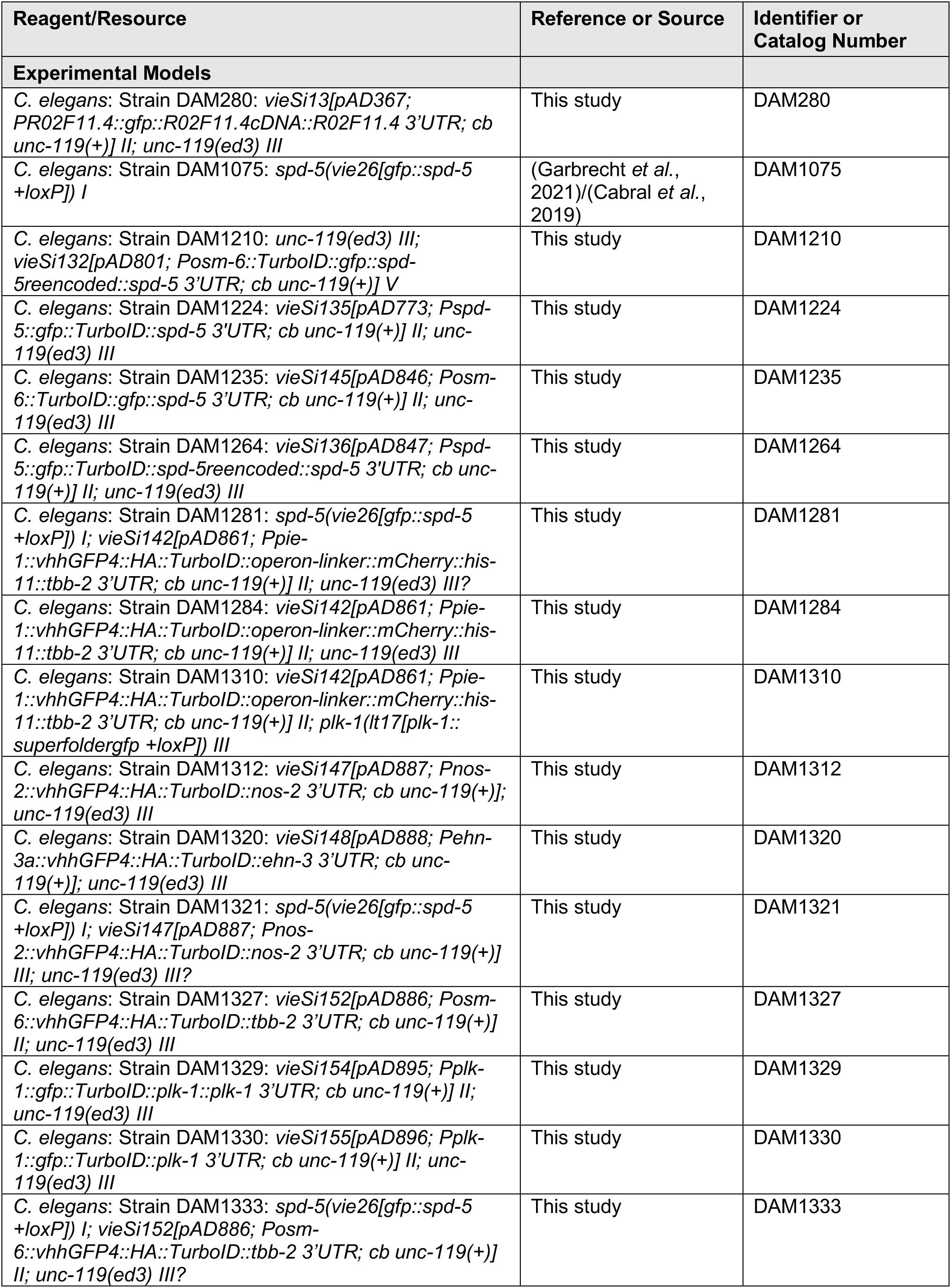
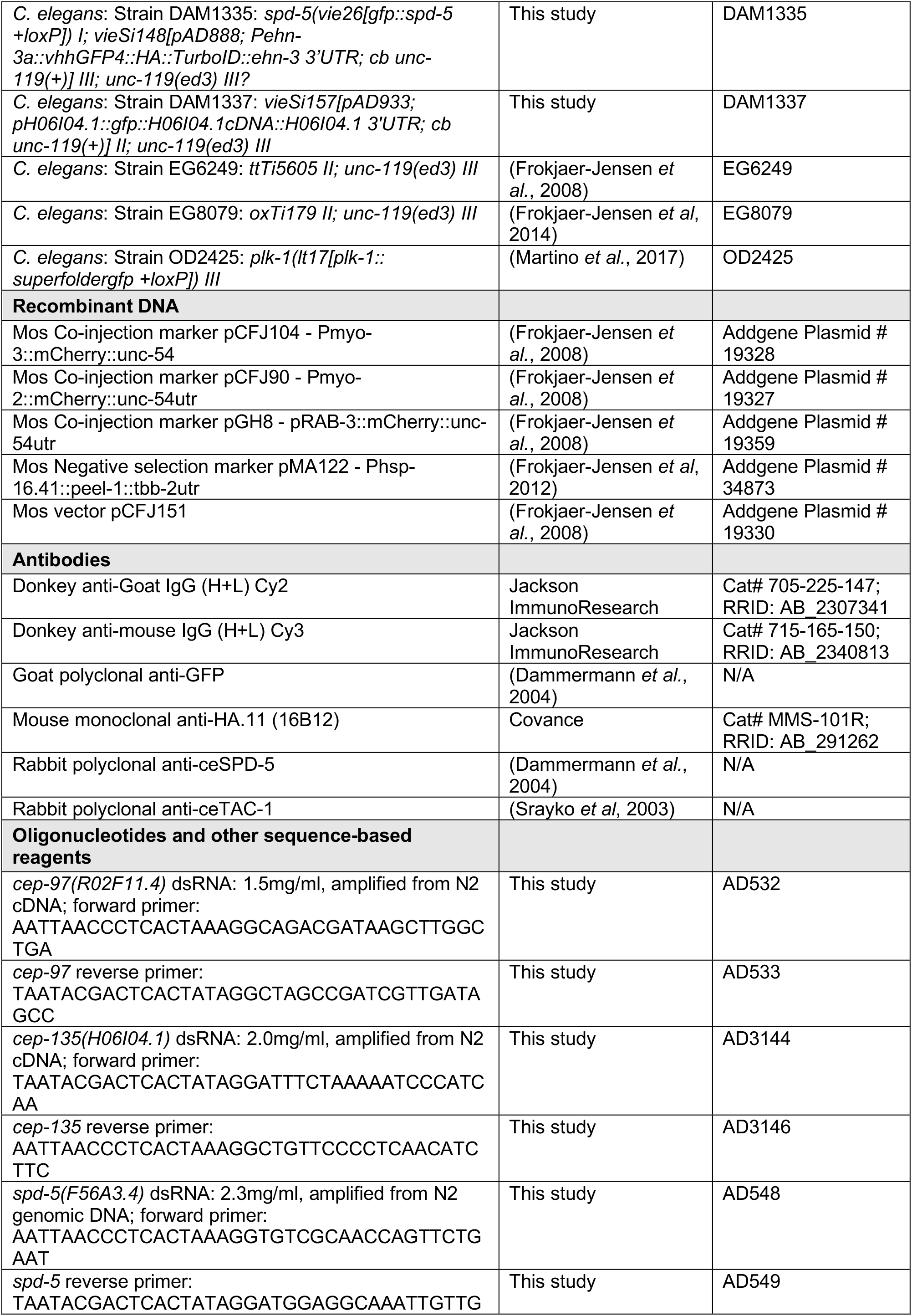
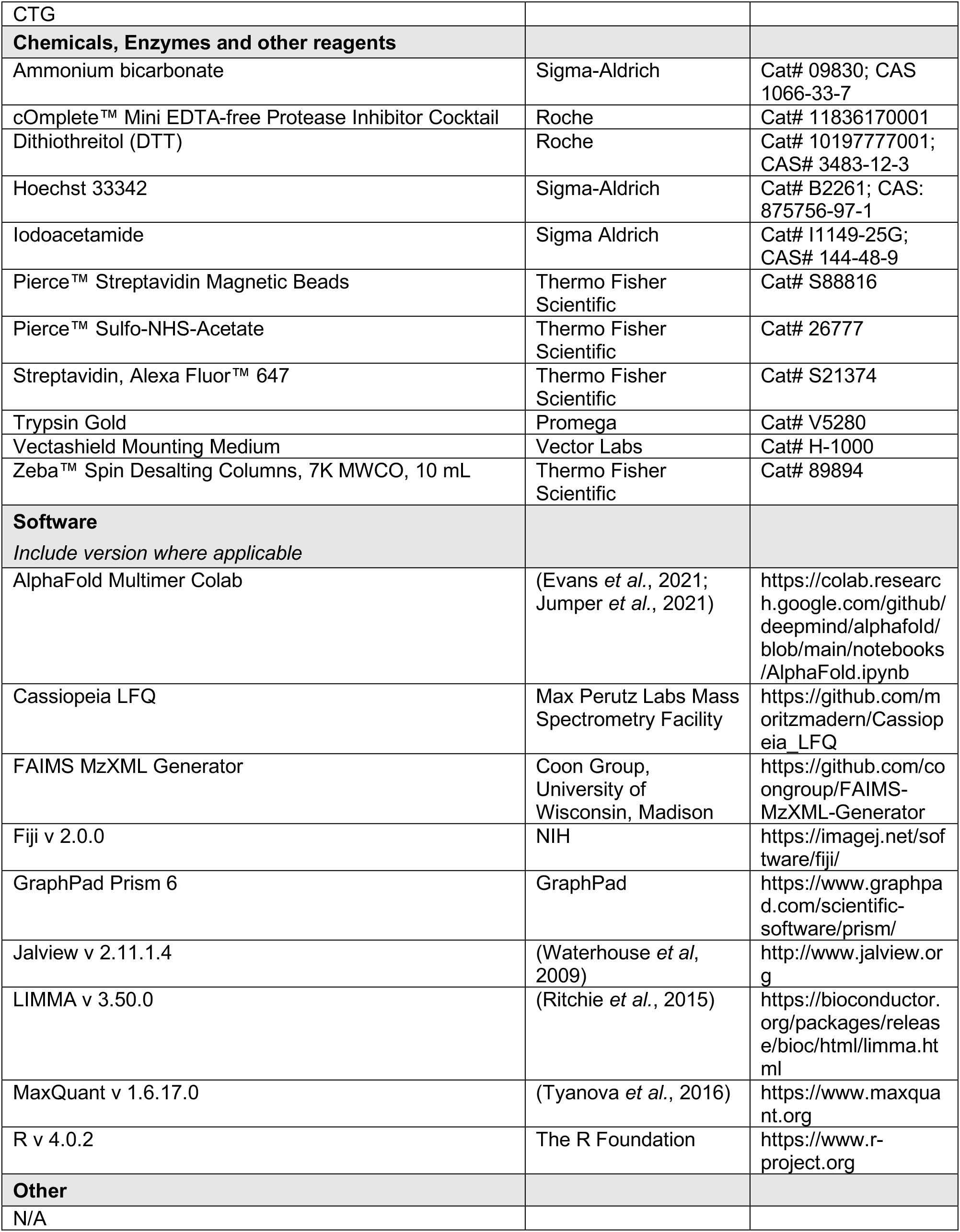

## ACKNOWLEDGEMENTS

We thank members of the Campbell and Dammermann labs for discussions, the *Caenorhabditis* Genetics Center, Luisa Cochella, Jessica Feldman and Karen Oegema for strains and reagents, Brooke Morriswood and Mario de Bono for sharing protocols and Markus Hartl of the Max Perutz Labs Mass Spectrometry facility and Josef Gotzmann and Thomas Peterbauer of the BioOptics facility for technical assistance. This work was supported by grants P30760-B28 and P34526-B from the Austrian Science Fund (FWF) to A.D..

## AUTHOR CONTRIBUTIONS

E.H. Conception and Design, Acquisition of Data, Analysis and Interpretation of Data, Drafting or Revising the Article; C.R-K. Conception and Design, Acquisition of Data, Analysis and Interpretation of Data; S.F. Analysis of 3D structure predictions; A.D. Conception and Design, Analysis and Interpretation of Data, Drafting or Revising the Article.

## SUPPLEMENTAL FIGURE LEGENDS

**Figure S1:**
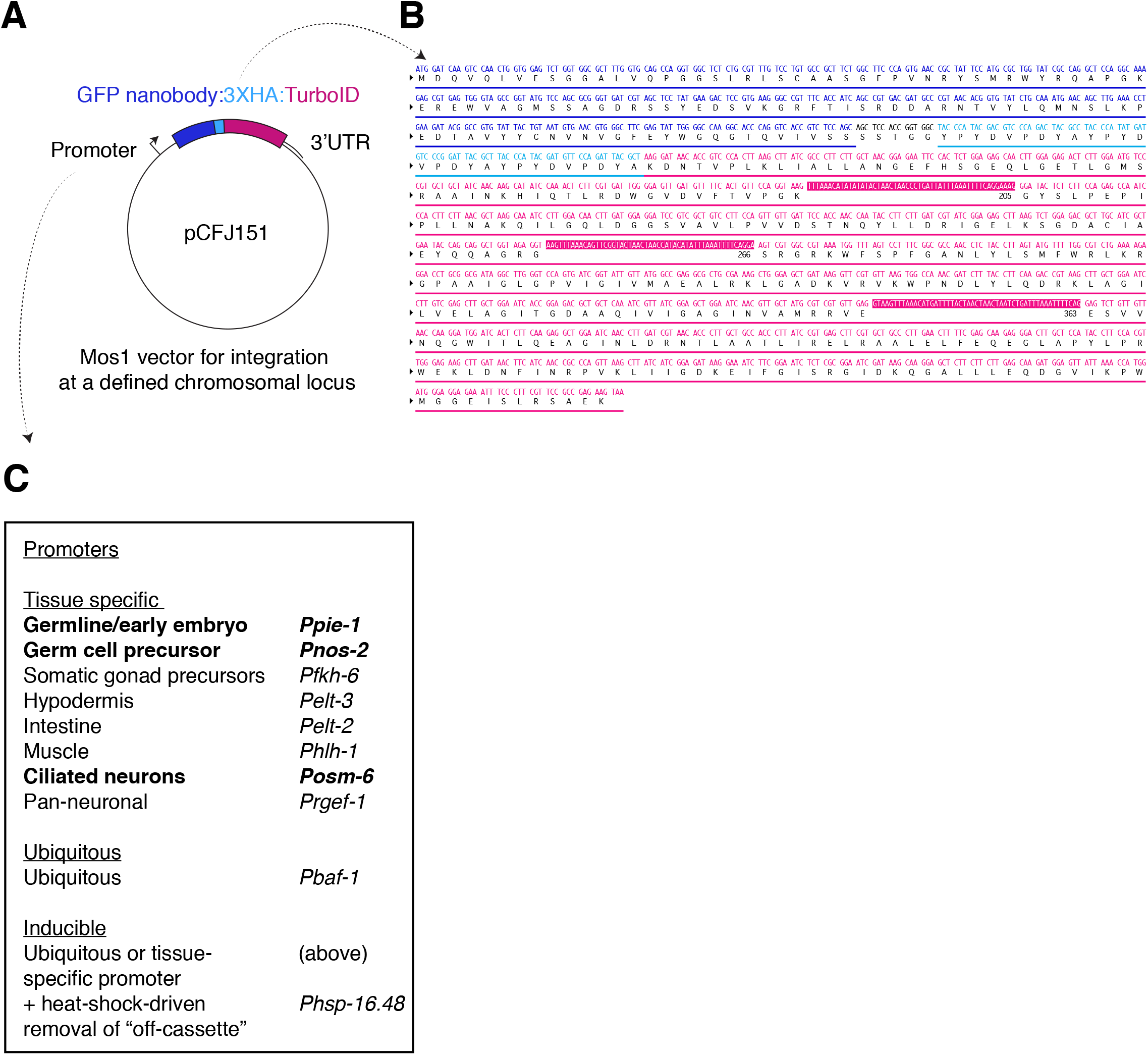
The GFP nanobody:TurboID tag. (**A**) Schematic of the indirect TurboID construct developed in this study. A GFP nanobody:HA:TurboID cassette is expressed under a ubiquitous or tissue/cell-specific promoter and 3’ regulatory sequences. The vector backbone is based on pCFJ151 for Mos1 transposon-mediated integration at a defined chromosomal locus (Frokjaer-Jensen *et al.*, 2008), sites for which have been established on all five autosomes in *C. elegans* (Frokjaer-Jensen *et al.*, 2014). (**B**) DNA sequence of the GFP nanobody:HA:TurboID cassette and conceptual translation. (**C**) Promoters for ubiquitous, tissue/cell-specific and inducible expression in *C. elegans*, selected for the low expression required for TurboID. Promoters in bold have been used and validated in this study. For inducible expression, a strategy such as using FLP-recombinase under the control of the *Phsp-16.48* heat shock promoter to excise a repressive “off-element” (Davis *et al*, 2008) avoids the variability and overexpression associated with the use of the heat-shock promoter alone and additionally confers the potential for tissue-specific inducible expression not explored in this study.

**Figure S2:**
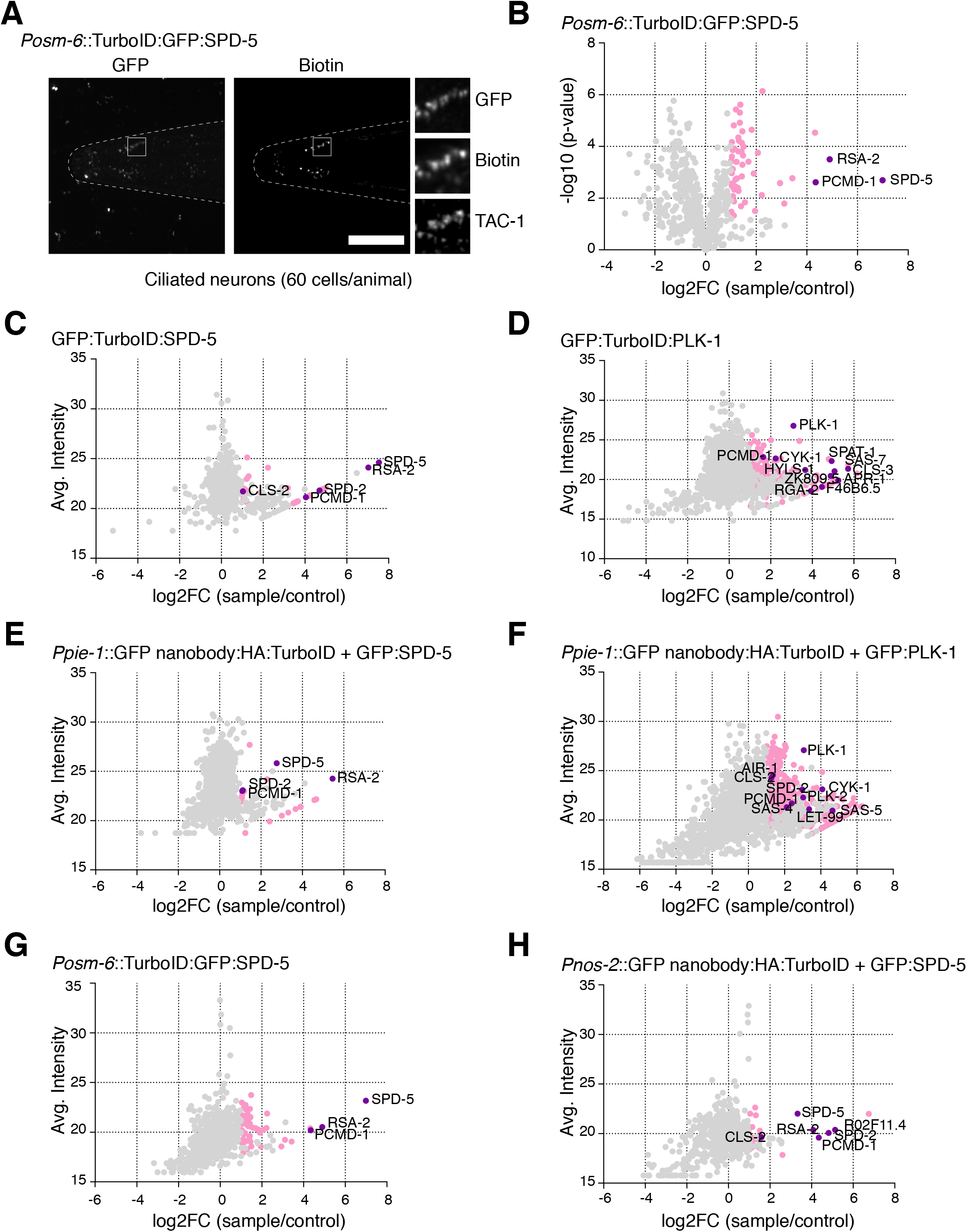
Tissue-specific labeling using direct TurboID and further data on TurboID experiments. (**A**) Direct TurboID applied to SPD-5 in ciliated neurons. Immunofluorescence micrograph of head of L1 larva from strain expressing TurboID:GFP:SPD-5 under the ciliated neuron-specific promoter *osm-6* stained for GFP, biotin (streptavidin) and TAC-1 as a PCM countermarker. Biotinylation signal is observed at the ciliary base coincident with GFP:SPD-5/TAC-1 signal. (**B**) Result of LC-MS/MS analysis for direct TurboID on SPD-5 in ciliated neurons. Volcano plot of −log10 p-values against log2 fold change (sample/control). Significantly enriched proteins (Log2 enrichment >1, p-value <0.05) are indicated in pink, with selected proteins highlighted. Comparison with indirect TurboID presented in Fig. 3C. See also Supplemental Table 1. (**C**-**H**) Volcano plots of log2 abundance (sample) against log2 fold change (sample/control) for direct (C, D) and indirect (E, F) TurboID on SPD-5 and PLK-1 in embryos, direct TurboID on SPD-5 in ciliated neurons (G) and indirect TurboID on SPD-5 in germ cell precursors (H). Significantly enriched proteins (Log2 enrichment >1, p-value <0.05) are indicated in pink, with selected proximity interactors highlighted. Note that interactors are present at much lower levels in the sample compared to endogenously biotinylated proteins, highlighting the importance of proper sample normalization. Scale bars in A is 10µm.

**Figure S3:**
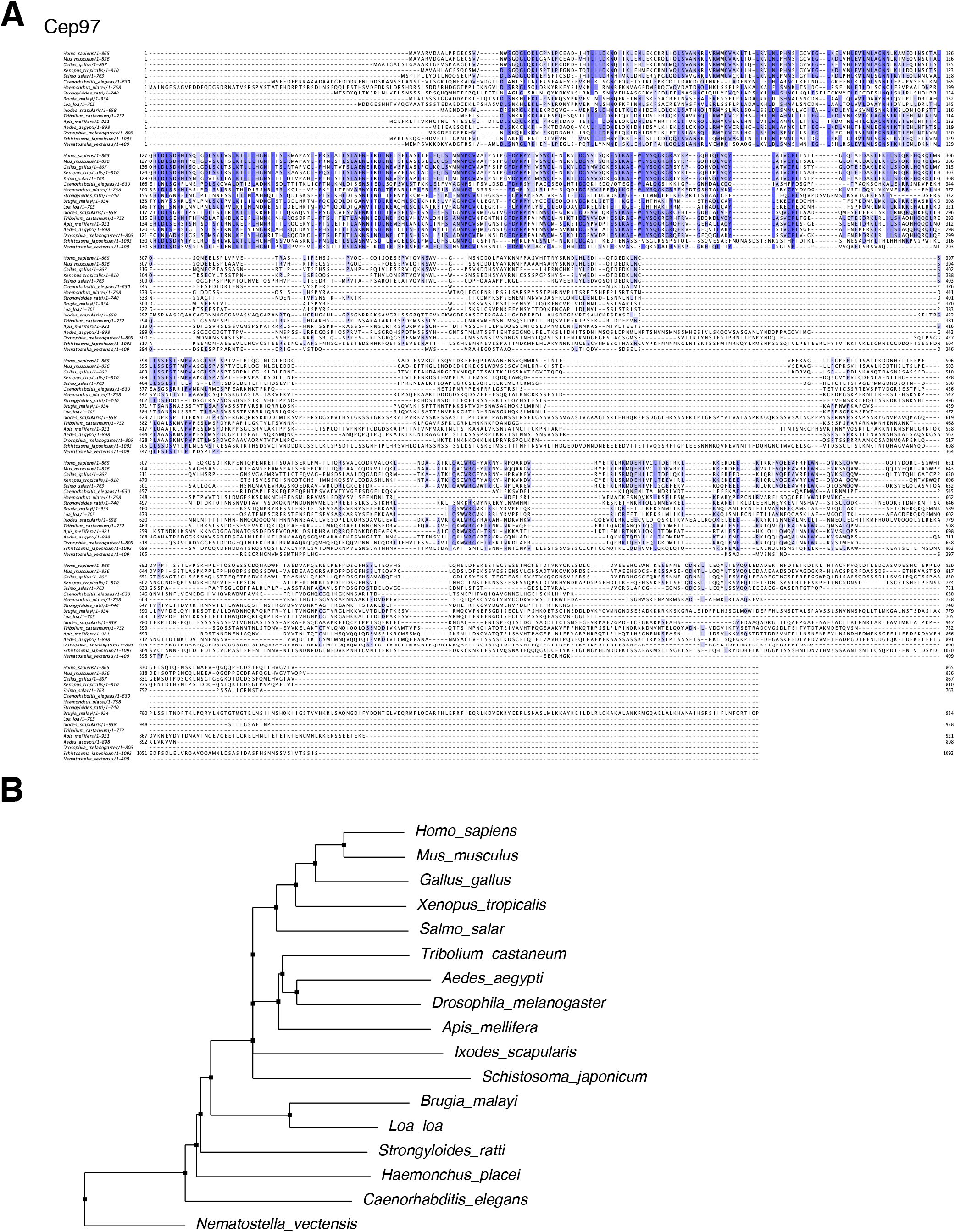
R02F11.4 as a putative ortholog of Cep97. **(A, B)** Multiple sequence alignment (A) and neighborhood joining phylogenetic tree (B) of selected Cep97 orthologs. Accession numbers are provided in Supplemental Table 2. Note that tree largely reflects pattern of evolutionary divergence.

**Figure S4:**
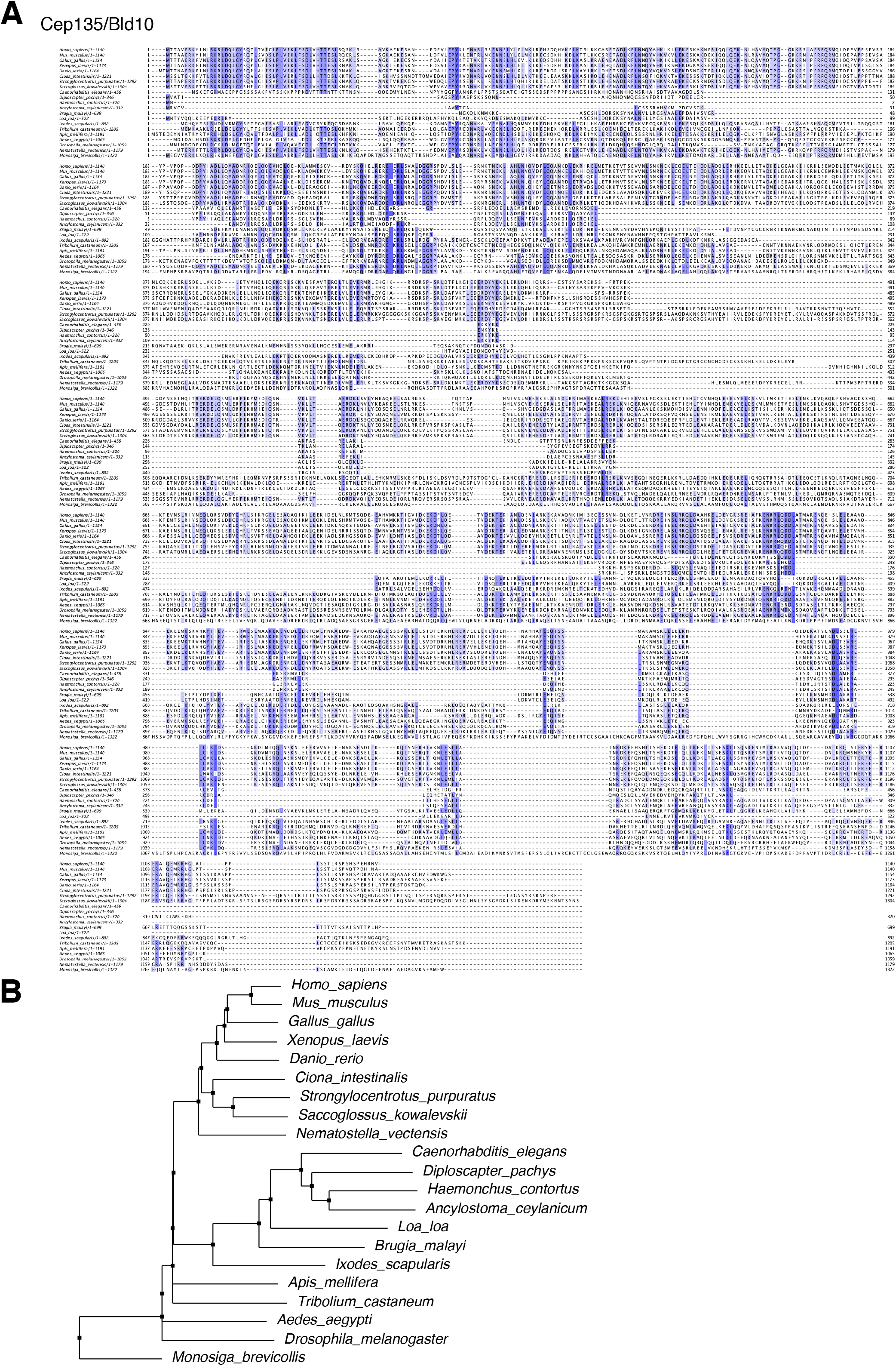
H06I04.1 as a putative ortholog of Cep135/Bld10. (**A**, **B**) Multiple sequence alignment (A) and neighborhood joining phylogenetic tree (B) of selected Cep135 orthologs. Accession numbers are provided in Supplemental Table 2. Note that tree largely reflects pattern of evolutionary divergence.

## SUPPLEMENTAL TABLES

**Table S1: Mass spectrometry data.** Complete list of proteins identified by mass spectrometry in each direct/indirect TurboID run, including corresponding controls.

**Table S2: List of Cep97/Cep135 orthologs.** GenBank accession numbers for Cep97/Cep135 orthologs presented in Figs 4, S3 and S4.

